# Targeted Molecular MRI of Colorectal Cancer by Antibody Functionalized Hyperpolarized Silicon Particles

**DOI:** 10.1101/2025.05.06.651536

**Authors:** Nicholas Whiting, Jingzhe Hu, Shivanand Pudakalakatti, Caitlin V. McCowan, Saleh Ramezani, Jennifer S. Davis, Niki Zacharias Millward, Brian J. Engel, Julie X. Liu, Klaramari Gellci, Hyeonglim Seo, Devin Brown, Jose S Enriquez, David G. Menter, Steven W. Millward, Seth T. Gammon, David Piwnica-Worms, Mary C. Farach-Carson, Daniel Carson, Pamela E. Constantinou, Pratip Bhattacharya

**Author notes:** **Address all correspondence to:** Pratip Bhattacharya, The University of Texas MD Anderson Cancer Center, Department of Cancer Systems Imaging, 1515 Holcombe Blvd., Houston, TX USA 77030. These authors contributed equally to this work as first author (Nicholas Whiting, Jingzhe Hu, Shivanand Pudakalakatti). Departments of Physics & Astronomy and Molecular & Cellular Biosciences; Rowan University, Glassboro NJ 08028. Department of Cancer Biology, The University of Kansas Medical Center, Kansas City, KS 66160. Sandia National Laboratories, Albuquerque, NM 87185. Undergraduate researcher (Klaramari Gellci, Devin Brown).

## Abstract

The development of non-invasive, non-ionizing sensitive molecular targeting approaches to detect colorectal cancer (CRC) lesions is warranted to improve high risk patient outcomes. Hyperpolarized silicon nanoparticles and microparticles are potentially well-suited to act as targeted molecular imaging agents because of their overall biocompatibility and long-lasting enhanced magnetic resonance imaging (MRI) signals. In this study, dynamic nuclear polarization was performed on silicon particles functionalized with an antibody to Mucin-1 (MUC1), a surface mucin glycoprotein aberrantly expressed in CRC. Antibody conjugation to the particle surface did not affect ^29^Si hyperpolarization characteristics. Similarly, conjugation and the dynamic nuclear polarization process did not adversely affect the affinity of the targeting antibody. *In vivo* MRI scans performed 10-15 minutes after luminal administration of targeted hyperpolarized particles into human MUC1-expressing orthotopic CRC mouse models showed that particles actively targeted tumor sites. These results were supported by chemical and biological controls and blocking experiments as well as correlative immunohistochemical analysis. These surface-functionalized silicon particles are under development as a platform technology that will allow non-invasive molecular targeting of CRC using hyperpolarized MRI.

**Single Sentence Summary:** Targeted molecular MR imaging of colorectal cancer by hyperpolarized silicon particles functionalized with mucin 1 antibody.

## Introduction

Silicon-based microparticles and nanoparticles are a focus of interest as targeted diagnostic and drug delivery vehicles. Their biocompatibility, biodegradability ^1^, and simple surface chemistry renders them amenable to drug loading and targeting ^2^. Many potential benefits exist for developing advanced biomedical imaging techniques that employ magnetic resonance imaging (MRI) because of its non-invasive, non-ionizing nature, as well as its capacity for high-resolution deep tissue imaging. Because ^29^Si (∼4.6% natural abundance) is MR-active (spin = ½), direct detection of silicon particles could prove useful for targeted molecular imaging because these particles provide positive contrast and background-free signal. To overcome the low sensitivity of ^29^Si MRI, researchers have explored a method of increasing the nuclear spin polarization of ^29^Si in microparticles and nanoparticles using solid-state dynamic nuclear polarization (DNP) ^3^, resulting in an increase in sensitivity of 4-5 orders of magnitude ^4^. DNP uses low temperatures and strong magnetic fields to generate a high electron spin polarization that is transferred to nearby nuclear spins through microwave-mediated dipolar interactions ^5^. This process occurs primarily on the surface of the particles and takes advantage of endogenous electronic defects at the Si-SiO_2_ interface of the naturally oxidized silicon surface ^6^, removing the need for exogenous radical species, which are frequently added for most small-molecule ^13^C or ^15^N DNP studies ^7^.Through nuclear spin diffusion ^8^, the polarization migrates to the core of the particle, where it is relatively well-protected from magnetization-depleting interactions with the immediate surroundings. Because of this, the relaxation rate (*T*_1_) of the hyperpolarized signal is exceedingly long (tens of minutes depending on particle size) and largely impervious to the chemical environment ^9^. This allows for a significantly longer imaging window for *in vivo* MRI studies than most other hyperpolarized contrast agents, which typically relax over the course of tens of seconds when administered to animal or human subjects ^10^. This extended imaging window mitigates one of the primary drawbacks of hyperpolarized contrast agents: limited imaging time driven by fast relaxation rates *in vivo* ^11^.

Previous work in this field conducted fundamental studies of silicon hyperpolarization ^3,4,8,12^ in particles ranging from the nanometer ^13^ to micron size scale, porous silicon nanoparticles ^14^, proof-of-concept *in vivo* imaging of untargeted silicon microparticles in mice ^9^, and the use of hyperpolarized silicon particles for MRI-guided tracking of angiocatheters ^15^. Another study ^16^ characterized the effects of coupling a thioaptamer to the particle surface to target E-selectin overexpression in ovarian cancer ^17^, demonstrating that particles conjugated to targeting moieties are suitable for *in vivo* imaging ^16^. Other hyperpolarized nanomaterials under investigation include ^13^C in nanodiamonds ^18,19^, phosphorus centers in doped silicon ^20^, silica nanoparticles ^21^, colloidal quantum dots ^22^, aluminosilicates ^23^, and carbonated hydroxyapatites ^24^.

The proof-of-concept work presented here demonstrates the utility of hyperpolarized silicon particles for targeted molecular imaging of CRC, the fourth leading cause of cancer-related deaths among adults in the United States ^25^. Despite the availability of preventive screening measures that include traditional colonoscopy, flexible sigmoidoscopy, fecal occult blood test, and fecal DNA analysis ^26–28^, many lesions remain undetected. Patient prognosis is typically favorable when CRC is detected sufficiently early (i.e., stages I or II) ^29^; unfortunately, a significant fraction of CRC is found after metastasis, when outcomes are less favorable ^25^. Traditional visual colonoscopies can fail to detect small or nonpolypoid colorectal neoplasms (i.e., flat lesions) and carry the risk of colonoscopic perforation ^30,31^. Although colonoscopic perforation is rare, it is associated with a high rate of morbidity and mortality ^31^. Additionally, CRC incidence is increasing significantly in younger populations and those with little surveillance ^32^. Development of advanced non-invasive imaging methods for early detection can improve clinical outcomes ^33^. Although implementation of virtual colonoscopy using computed tomography (CT) ^34^ or MRI ^35,36^ has increased, these techniques suffer from drawbacks that include difficulty in detecting small (<10 mm) tumors and lesions ^37^, susceptibility distortions ^38^, radiation exposure ^39^, false positives from residual fecal matter ^40^, bowel preparation, and reliance on non-tumor-specific contrast media ^41,42^. Given the increase in CRC incidence amongst individuals younger than 55 years ^32^, repeated CT-based virtual colonoscopies may prove impractical because of cumulative radiation exposure.

The colon lumen enables the use of probes that target epitopes on the epithelial cell surface without a need or vascular delivery. Development of a positive contrast agent to selectively target CRC molecular biomarkers using non-invasive, non-radiative MRI could enhance the impact of virtual colonoscopies in patient care. Mucins are a class of glycoproteins expressed by most luminal epithelial cells. Mucins provide a physical barrier to protect and lubricate cells and simultaneously link to cell signaling pathways ^43^. MUC1 possesses a heavily glycosylated ectodomain that extends 200-500 nm from the cell surface that actively binds pathogens and prevents them from penetrating the epithelium ^44,45^. Overexpression of MUC1 and aberrant glycosylation are typical in most epithelial cancers ^46^, including CRC, where it has been positively correlated with CRC proliferation and negative outcomes ^47^. MUC1 overexpression blocks chemotherapeutic attack and blunts immune system response while enhancing proliferative and anti-apoptotic signaling pathways ^43^. Surface MUC1 has been leveraged for targeted imaging and therapy ^48–50^ including antibody targeting of shed MUC1 in serum ^51^, immune response therapy to MUC1 overexpression ^52^, and small molecule inhibition therapy that targets the MUC1 cytoplasmic tail ^53^.

The large glycosylated ectodomain of MUC1, extending 500 nm from the cell surface, can be recognized by targeted nanoparticles, such as quantum dots ^54^, gold nanoparticles ^55^, and polymer nanoparticles ^56^. Particle conjugation to glycoform-specific antibodies, which target the tandem repeat region of the MUC1 ectodomain, allows for multiple potential binding sites, further amplifying the signal ^55^ and increasing targeted imaging sensitivity. Here, the human MUC1 (huMUC1)-specific *214D4* antibody ^57,58^ was coupled to the surface of silicon microparticles (average mean diameter ∼2 μm) to target MUC1 using hyperpolarized ^29^Si MRI. While microscale silicon particles typically present challenges for preclinical molecular imaging because of their large size and decreased mobility ^16^, targeting MUC1 in CRC leverages a unique set of favorable conditions. MUC1 ectodomain is directly available to agents delivered intraluminally, in this case via enema, instead of relying on intravenous injections. Direct delivery circumvents particle blockage of block blood vessels and mitigates the time constraints from natural decay of hyperpolarized MR signal. Direct delivery protects the functionalized antibodies on the surface of silicon particles from degradation and minimizes exposure of other organs to both the antibody and particles.

In this work, we present data demonstrating that functionalized silicon particles can target MUC1 in CRC using hyperpolarized MRI. *In vivo* studies using huMUC1-expressing CRC mouse models (subcutaneous and orthotopic) demonstrated that the hyperpolarized ^29^Si signal allows imaging after particle administration, and the bound particles allow for highly selective and specific tumor discrimination using ^29^Si MRI.

## Results

### MR virtual colonoscopy using hyperpolarized ^29^Si

Hyperpolarized PEGylated silicon particles (125 mg of polydisperse ∼2 μm diameter) were dispersed in phosphate-buffered saline solution (PBS) and administered to the lower intestinal tract of a live mouse through the rectum (in a procedure similar to an enema). The ^29^Si MRI scan was performed 5 min after particle administration, followed by the ^1^H anatomical scan. The co-registered images showed an even distribution of particles from the rectum to the cecum (Fig. 1, Fig. S1). While the mouse had no CRC tumors and the particles were not specifically targeted, the clear visualization of the lower intestinal tract using hyperpolarized ^29^Si particles (over the timescale of several minutes) mimicked the appearance of a conventional diagnostic barium enema and served as justification for further development of targeted hyperpolarized ^29^Si imaging of the bowel. Following the imaging experiment, the mouse was euthanized and the large intestine was excised. The solution of black particles could be clearly visualized (through the intestinal wall) inside the same section of intestinal tract that provided the ^29^Si signal in the *in vivo* image (Fig. S4).

**Figure 1:**
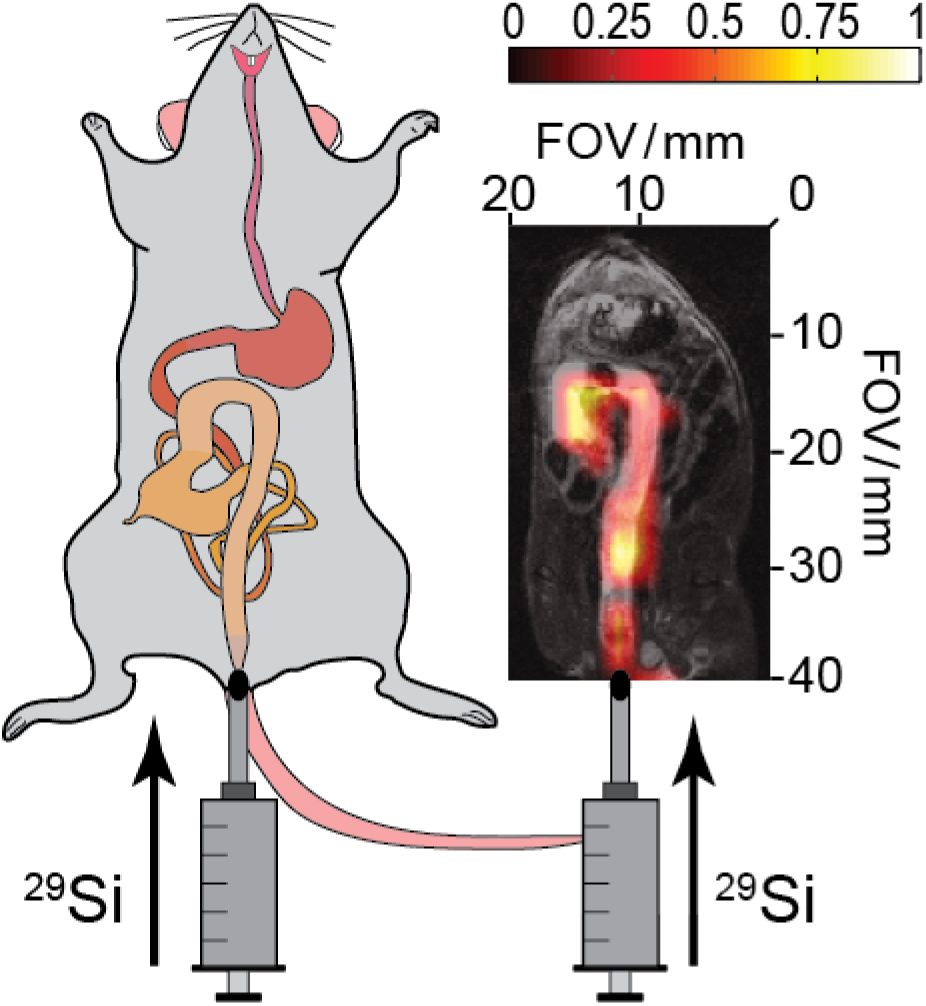
Proof-of-concept for silicon particle colonoscopy. (*Left*) Schematic demonstrating the rectal injection of silicon particles into the lower intestinal tract. (*Right*) Rectal injection of 125 mg of PEGylated silicon microparticles (in 500 μL PBS) into a normal mouse; ^29^Si image taken 5 minutes post-injection. ^29^Si MRI (*color*) overlaid on ^1^H MRI (*greyscale*) showing silicon particles occupying the intestines from the rectum to the cecum. Imaging parameters are available in the **Supplemental Section 2**, as is an uncropped version of the ^29^Si/^1^H overlay image (**Supplemental Figure S1**).

### Effects of antibody functionalization on ^29^Si DNP

The *214D4* antibody was chosen for targeting huMUC1 because of its affinity and specificity toward the glycosylated ectodomain of the transmembrane glycoprotein ^55,57^. Conjugation of an antibody may adversely affect the hyperpolarization characteristics of the silicon particles via alterations in the position and number of endogenous electronic defects due to changes in the surface chemistry. However, whether the hyperpolarization characteristics would be affected in this case was unknown at the outset. Fig. 2 compares the hyperpolarized ^29^Si signal in unfunctionalized vs. *214D4*-functionalized microparticles, showing nearly identical initial ^29^Si MR signals and highly similar *T*_1_ decay rates (*T*_1_ ∼19 min). This capacity to hyperpolarize the antibody-functionalized silicon microparticles as efficiently as native particles demonstrates their viability for use as targeted imaging agents for ^29^Si MRI. Furthermore, useful hyperpolarized ^29^Si MR spectra were achieved at room temperature up to 40 min post-DNP, which suggests that imaging studies with this system could be carried out for ten times longer than conventional hyperpolarized MR studies.

**Figure 2:**
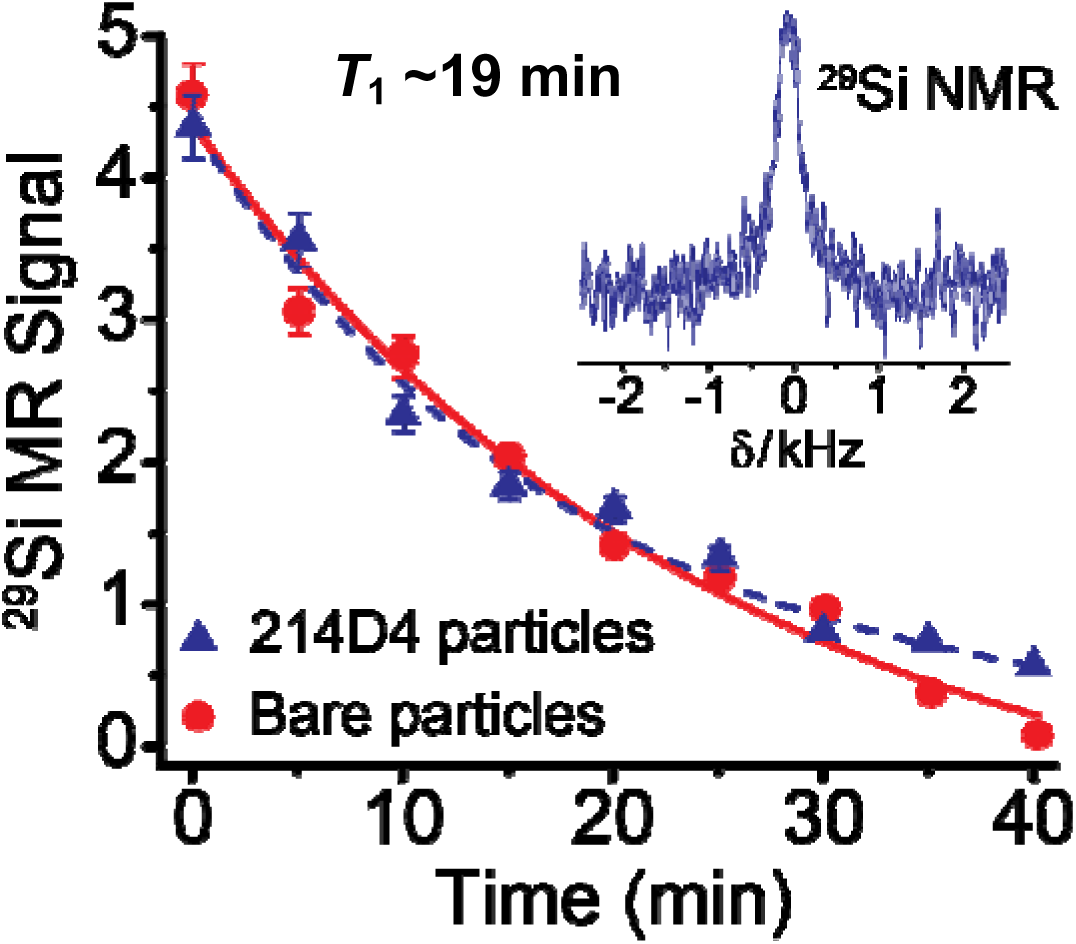
Antibody functionalization does not impair ^29^Si DNP. ^29^Si MR signal decay curves for 2 μm silicon microparticles that are both: conjugated to *214D4* antibodies to target MUC1 (*triangles*), and unfunctionalized (*circles*). Samples possess similar initial ^29^Si signal and decay constant (HP *T*_1_ ∼19 min for both samples). Targeted sample consisted of 60 mg silicon particles coupled to *214D4* antibody (see text) in 150 μL HEPES buffer and polarized for 16.5 hrs. Unfunctionalized sample consisted of 61 mg silicon particles in 150 μL HEPES buffer and polarized for 18 hrs. Error bars indicate variations in scanner sensitivity through comparison with a silicon reference standard immediately prior to data collection. *Inset*: ^29^Si MR spectrum from *214D4* functionalized silicon microparticle sample immediately after placement in 7 T Bruker MR scanner (*T*=0). The spectrum is taken with 10^0^ pulse.

### Effects of ^29^Si DNP on antibody functionality

After demonstrating that the addition of the *214D4* antibody had no effect on the hyperpolarization characteristics of the particles, it was important to verify that the harsh conditions of DNP did not affect the capacity of the antibody to target the ectodomain of huMUC1. While the hyperpolarized ^29^Si signal lasts for tens of minutes, it is not nearly long enough to allow antibody functionalization steps to be performed after DNP. Therefore, the hyperpolarization process must take place on pre-conjugated particles (Fig. 3A). During DNP, the functionalized particles were exposed to temperatures of ∼3 K and irradiated with an 80.9 GHz millimeter wave source (100 mW) for up to 18 h, potentially altering the antibody’s structure and affinity. To validate the integrity of the antibody after this process, a small aliquot of *214D4* antibody solution (in a buffer; not conjugated to silicon particles) was exposed to the DNP conditions and then compared to a non-DNP sample by western blot (Fig. 3B). We found that the *214D4* antibody withstood the conditions of DNP without losing its capacity to engage the huMUC1 ectodomain, as quantified by densitometry (Fig. 3C). Furthermore, *in vitro* studies were conducted on particles exposed or not exposed to DNP to identify any noticeable differences in the capacity of the particles to bind to a MUC1-expressing human CRC cell line (Fig. 3D). The *214D4*-functionalized particles bound to huMUC1-expressing cells (HT29-MTX-E12) regardless of whether they had been exposed to DNP. Particles functionalized with only polyethylene glycol (PEG; i.e., without antibody) did not bind to huMUC1-expressing cells (chemical control), and *214D4*-functionalized particles did not bind to non–MUC1-expressing HS-5 cells (biological control). This demonstrates that the DNP process did not significantly affect the target-binding affinity of the *214D4* antibody nor the integrity of the coupling scheme that links the antibody to the particles. Furthermore, the particle binding was specific and targeted.

**Figure 3:**
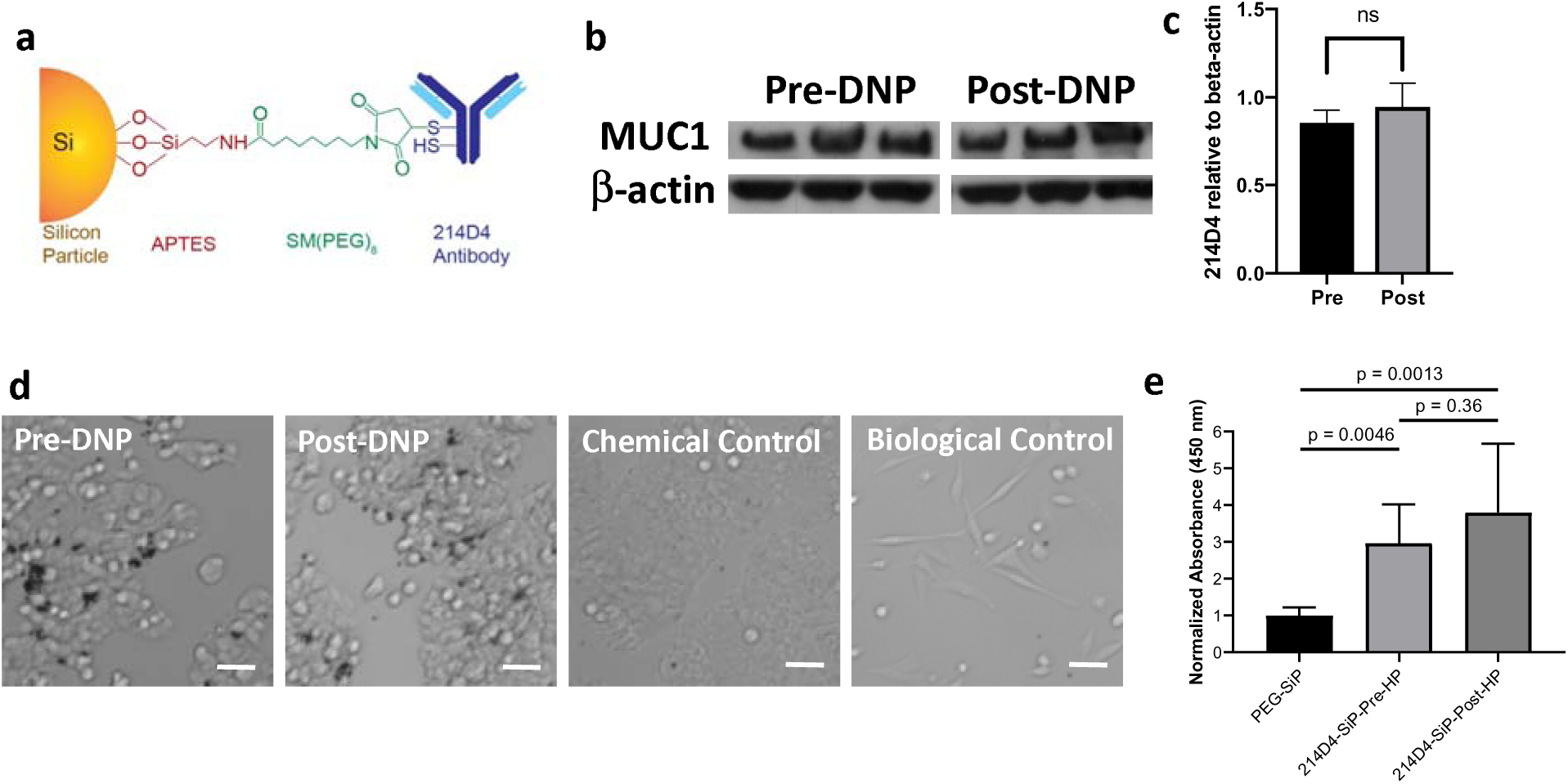
^29^Si DNP does not impair antibody recognition. The process of DNP does not affect the targeting sensitivity or specificity of MUC1-directed antibodies (*214D4*), as shown by three different methods. **a** Schematic demonstrating the coupling of *214D4* antibody to the silicon particles via APTES and heterobifunctional PEG linkers. **b** Western blot of lysate from HT29-MTX-E12 cells probed MUC1 antibody (pre- and post-DNP treatment) retained its full ability to bind the MUC1 epitope. **c** Densitometric analysis of western blot shows no reduction in the MUC1 antibody binding affinity in the antibody subjected to DNP conditions. **d** *In vitro* binding assays demonstrate that silicon particles (black dots) conjugated to *214D4* antibody successfully bind to HT29-MTX-E12 cells both before and after DNP. Non-targeted silicon particles do not bind to HT29-MTX-E12 cells (chemical control), and *214D4*-targeted particles do not bind to the HS-5 cell line (MUC1^−^ biological control). Scale bars indicate 100 µm. **e** Modified ELISA shows that DNP treatment of *214D4*-conjugated silicon microparticles does not affect binding of recombinant MUC1 ectodomain.

The retention of antibody functionality after the DNP process was validated by a third method. A modified enzyme-linked immunosorbent assay (ELISA) with silicon microparticles was performed with a recombinant biotinylated MUC1 ectodomain fragment (Fig. S11). PEGylated *214D4* conjugated pre-DNP or *214D4* conjugated post-DNP was incubated with a biotinylated huMUC1 ectodomain and then with streptavidin–horseradish peroxidase (HRP) and developed with a 3,3′,5,5′-tetramethylbenzidine (TMB) substrate. DNP caused no significant change in signal (Fig. 3E). These data demonstrate that *214D4* can withstand the conditions of DNP without loss of its capacity to target the MUC1 ectodomain.

### Prolonged ^29^Si imaging in subcutaneous CRC mouse model

Initial mouse models of CRC were generated by subcutaneous injections of huMUC1-expressing CRC cells. Because of the high viscosity of the silicon microparticle solution, tail vein injections were not feasible ^16^. Therefore, the hyperpolarized particles were injected directly into the tumor. A strong ^29^Si signal remained at the injection site even 20 min after injection (Fig. 4). While deposition *in situ* is not considered sufficient to demonstrate molecular targeting, the ability to acquire MR images of targeted particles in a tumor *in vivo* over long time durations directly demonstrates the feasibility of this contrast method. An additional example of ^29^Si imaging in a subcutaneous tumor model is presented in Fig. S3.

**Figure 4:**
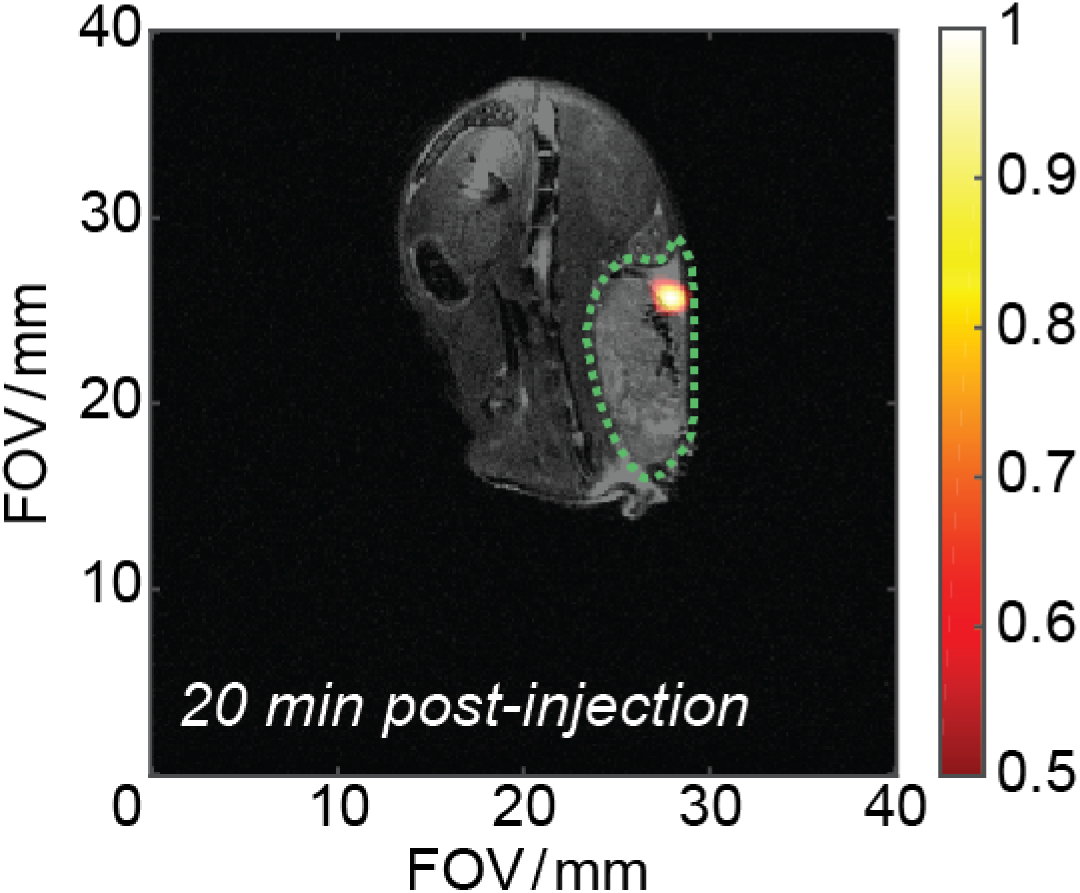
Long lasting HP ^29^Si MR signal inside tumor volume. 60 mg of *214D4*-functionalized 2 μm silicon microparticles (in 130 μL PBS; *T*_pol_ ∼18 hr) directly injected into tumor volume of a MUC1-expressing subcutaneous HT29-MTX-E12 CRC mouse model. Image taken 20 min after intratumoral injection. Silicon image (*color*) overlaid with ^1^H anatomical image (*greyscale*) for co-registration; tumor volume outlined with dotted green line. Imaging parameters are available in the **Supplemental Section 2**, as is a similar study (**Supplemental Figure S2**) with the image taken 5 minutes after injection into a different mouse.

### Targeted imaging in an orthotopic CRC mouse model

To demonstrate the targeting capacity *in vivo* of the *214D4* particles, a transgenic huMUC1-expressing orthotopic CRC mouse model was developed that produced tumors in the lower intestinal tract between the rectum and the cecum (Fig. S6). It should be noted that although wild-type mice express MUC1, the *214D4* antibody is specific to the ectodomain of huMUC1 (Fig. S6). Tumor volumes ranged from 14-66 mm^3^, and tumor presence and location were validated by optical colonoscopy (Supplemental Video SV1) prior to the MRI experiments. The hyperpolarized MUC1-targeted microparticles were administered to the mice via the rectum, and the ^29^Si signal was acquired after a 10- to 15-min wait period (Fig. 5). Although previous experiments (including Fig. 1) indicated that the silicon particle solution effectively coated the intestines from the rectum to the cecum, an initial ^29^Si MR spectrum (α=10°) was acquired immediately after particle administration to confirm the integrity of the available hyperpolarized ^29^Si MR signal to be used in the imaging experiment. In these studies, concentrated ^29^Si MR signal was present in the colonoscopy-validated tumors (Fig. 5, n=3).

**Figure 5:**
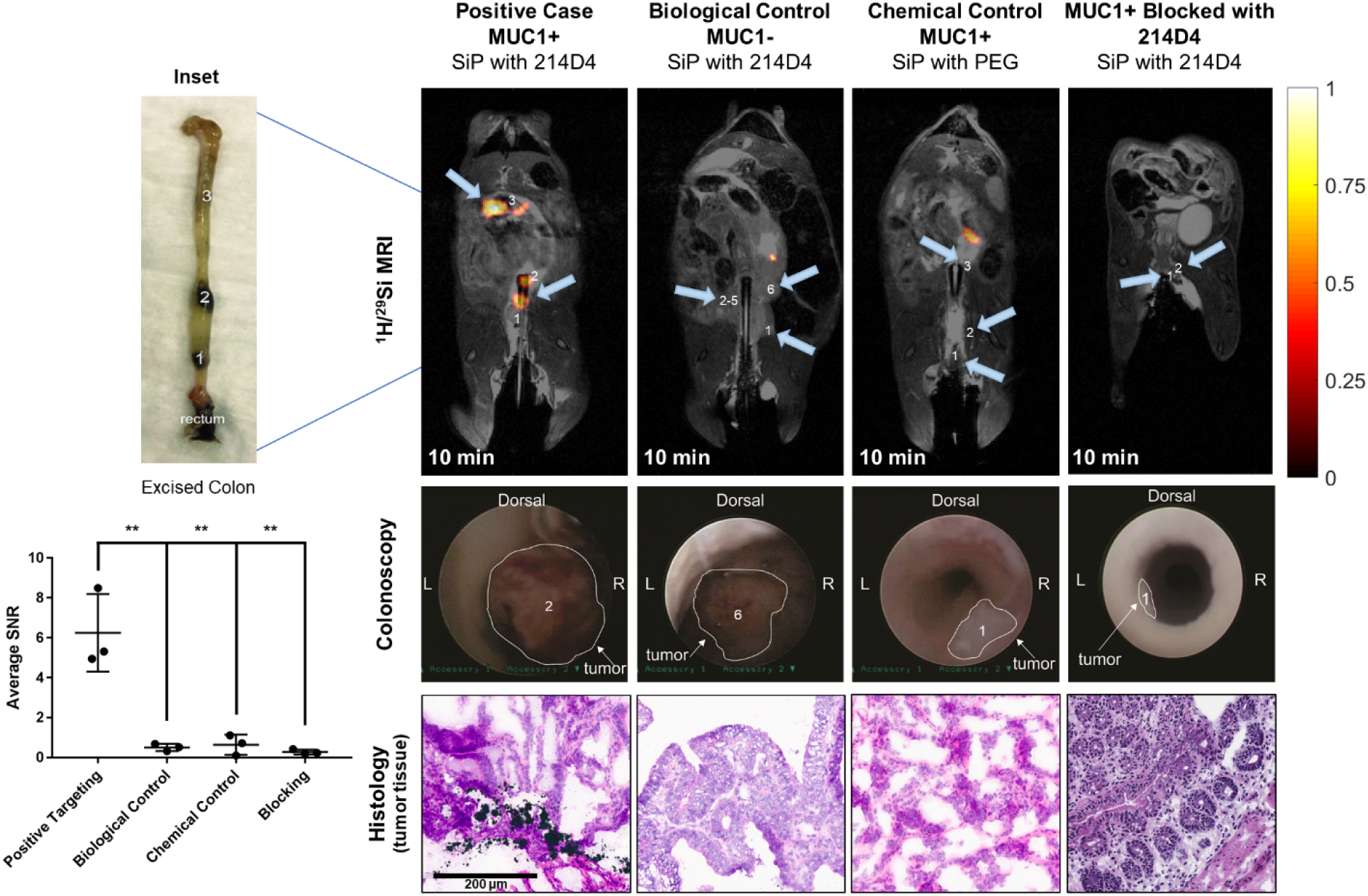
Targeted molecular imaging of CRC. Functionalized silicon microparticles administered to a humanized CRC mouse model (MUC1^+^). 45 mg of 2 μm HP SiPs in ∼300 μL PBS, *T*_pol_ ∼18 hr, administered through the rectum of mice. **Upper panel:** ^29^Si signal hyperpolarized images (*color*) overlaid with ^1^H anatomical image (*greyscale*) for co-registration; Co-registered ^29^Si/^1^H MRI scans taken 10 min after particle administration; all experiments repeated (N=3). ***Far-left column*** denotes silicon particles functionalized with *214D4* antibody administered to MUC1+ mice with CRC tumors. ***Left-middle column*** denotes a biological control study using *214D4*-functionalized particles administered to a mouse with a non-MUC1-expressing CRC tumor (MUC1-). ***Right-middle column*** denotes a chemical control study using PEGylated (non-targeted) particles in the MUC1+ CRC mouse model. These two middle columns represent biological and chemical controls to establish targeting efficacy. ***Far-right column*** denotes a control blocking study using *214D4* antibody functionalized particles administered to pre-blocked MUC1+ mouse with CRC tumors. Numbers and blue arrows show the number and location of tumors in the mouse, respectively. **Middle panel:** representative images of the matching colonoscopy from a Karl Storz Coloview veterinary mini-endoscopic system. Imaging parameters are available in the **Supplemental Section 2**. **Lower panel**: Histology of the excised tumors from the corresponding animals correlates with imaging studies. Numerical assignments of tumors visualized by ^29^Si MRI and endoscopy respectively are shown. **Upper-left inset:** Photograph of the excised colon from the positive control, showing the silicon particles binding to the tumor sites. **Bottom-left inset:** Comparative statistical analysis of molecular targeting ability of *214D4* antibody functionalized hyperpolarized silicon particles through signal to noise (SNR) analysis in MUC1 + animals (N=3), biological (N=3) and chemical controls (N=3) and blocking trials (N=3); ** denotes *p* < 0.01. Details of calculation are outlined in **Supplemental Section 2**.

In control studies, PEGylated (non-targeted) silicon particles were administered to a mouse littermate of the same genotype, *214D4*-functionalized silicon microparticles were administered to non–MUC1-expressing orthotopic CRC mice, and *214D4*-functionalized silicon microparticles were administered to huMUC1-expressing orthotopic CRC mice in which huMUC1 was pre-blocked with fluorescently labeled unconjugated *214D4* (*214D5*/Cy5; Fig. S8). This set of control experiments was repeated (n=3) per condition to verify reproducibility; all trials showed similar results (Figs. S7, S9). In all control experiments, the concentrated ^29^Si MR signal was absent in the tumor sites. Instead, in several experiments a faint background of ^29^Si MR signal was observed at non-localized positions within the intestines, which is to be expected in the absence of a “wash-out” step. The signal was faint because of the diluted nature of the administered particles (i.e., fewer particles per pixel) when they are not actively targeting a tumor.

Immediately following each experiment, mice were euthanized, and the large intestines (from the rectum to the cecum) were removed, flushed with PBS, preserved in optimal cutting temperature (OCT) compound, and subjected to tissue analysis. After sectioning, washing, and staining, optical microscopy (Fig. 5) clearly indicated that the targeted silicon particles adhered to the cell surface of the MUC1-expressing tumor cells. This adhesion was not observed in the chemical or biological controls from any of the mice (Fig. S10). This analysis confirmed that the *214D4*-functionalized silicon particles actively targeted and bound the MUC1-expressing tumor cells, confirming the ^29^Si MRI results.

## Discussion

This work established the utility of hyperpolarized silicon particles for targeted molecular imaging of CRC tumors using non-invasive MRI. We demonstrated that rectal administration of hyperpolarized silicon particles via enema can discern the lower intestinal tract of a mouse from the rectum to the cecum using ^29^Si MRI. In the case of a normal mouse and non-targeted silicon particles, the contours of the bowel were readily visible in positive contrast for several minutes after particle administration (similar in function to a barium enema). Hyperpolarization of native silicon microparticles has been previously reported ^9^, as have hyperpolarized nanoscale silicon particles functionalized with small molecules ^12^ or aptamers ^16^. However, there have been no previous reports on the effects of antibody functionalization on the surface of silicon microparticles. Because the particles utilize endogenous surface electronic defects to drive the DNP process (i.e., they are not reliant on exogenous radicals), changes to their surface chemistry could affect their capacity to generate enhanced ^29^Si MR signals. We demonstrated that there is essentially no difference in the initial level of ^29^Si MR signal or the room temperature *T*_1_ rate for particles conjugated with or without the *214D4* antibody (Fig. 2). The nearly identical initial ^29^Si MR signals between samples indicate that the hyperpolarization efficiency and overall nuclear spin polarization values are unaffected by the surface chemistries used in this study. Furthermore, the long *T*_1_ (∼19 min for both samples) allows useful ^29^Si signal to be observed for 40 min after placement of the sample into the MR scanner. These findings suggest that these antibody-functionalized silicon particles are sufficiently hyperpolarized, clearing an important hurdle in the development of these particles for targeted molecular imaging.

It was equally important to verify that the potentially non-biological conditions of DNP (irradiation with 100 mW of 80.90 GHz millimeter waves at ∼3 K for as long as 18 h) did not negatively affect the targeting efficiency of conjugated antibody. When the polydispersible antibody-functionalized silicon microparticles were hyperpolarized using DNP and compared to non-DNP samples in an *in vitro* binding assay, the two groups showed similar binding to huMUC1-expressing HT29-MTX-E12 cells and the purified ectodomain of MUC1 (Fig. 3). To our knowledge, this is the first confirmation that the conditions of DNP do not affect the targeting function of an antibody and bodes well for sustainable development of hyperpolarized silicon particles for molecular imaging using other antibody-based targeting reagents. As was expected, neither sample demonstrated appreciable non-specific binding to control cells that did not express MUC1.

Initial *in vivo* imaging studies using *214D4*-functionalized silicon microparticles were carried out using mice with subcutaneous (flank) MUC1-expressing CRC tumors. The primary goal of these experiments was to confirm that the hyperpolarized ^29^Si signal could still be observed *in vivo* using the targeted particles and to establish a relevant time window for imaging studies. *In vivo* imaging using hyperpolarized silicon particles can be challenging compared to MR spectroscopy because of spatial delocalization of the particles, resulting in less signal per pixel/voxel. Because of previous difficulties in employing tail vein injections to administer micron-size silicon particles ^16^, the particles were instead administered by direct (intratumoral) injection. A hyperpolarized ^29^Si signal was achieved for at least 20 min after the injection, with the signal remaining within the tumor volume. An additional study in another mouse with signal acquisition after 5 min (Fig. S3) showed a similar signal intensity at the injection site, with a slightly broadened spread of particles. While this particular trial cannot be considered true molecular targeting, it helps to establish a relevant timeline for the acquisition of hyperpolarized ^29^Si signal *in vivo* and confirms that hyperpolarized ^29^Si MR signal from the *214D4*-functionalized particles can be observed within a tumor environment.

Our transgenic orthotopic mouse model that spontaneously produces huMUC1-expressing CRC in the lower intestinal tract allowed for administration of the hyperpolarized silicon particles through the rectum, where they could directly access the glycosylated ectodomain of the MUC1 expressed on the CRC tumor surface. Given our success in acquiring hyperpolarized ^29^Si MR signal concentrated at a tumor site within 20 min, wait periods of 10-15 min were used to allow the particles time to bind before signal acquisition. This pre-acquisition waiting period served as a temporal signal wash out. Because unbound particles were not physically removed or flushed before signal acquisition, there needs to be a method of differentiating hyperpolarized ^29^Si signal from particles bound to the tumor from the signal from particles not bound. Particles that are bound to the tumor were concentrated within a small number of pixels, increasing signal intensity at the tumor site. In contrast, unbound particles were distributed throughout the intestines, resulting in a lower ^29^Si signal per pixel. The chosen wait time allowed sufficient magnetization loss due to hyperpolarized *T*_1_ processes so that the lower concentrations of particles that were unbound produced minimal signal compared to particles that were concentrated at the tumor, but still within the time envelope for high signal-to-noise molecular targeting. Future experiments will test the addition of a physical washout step (i.e., saline enema flush) between particle administration and MRI.

These studies demonstrated that *214D4*-functionalized microparticles provided concentrated ^29^Si signal at the huMUC1-expressing tumor but not in the three controls (targeted particles in a non– MUC1-expressing mouse, untargeted particles in a huMUC1-expressing transgenic mouse, and targeted particles in a pre-blocked huMUC1-expressing transgenic mouse). Furthermore, post-imaging histologic analysis of excised intestines confirmed that only *214D4*-functionalized particles bound to the MUC1-expressing tumors. No evidence of particle binding was observed in the *in vivo* chemical and biological controls or after pre-blocking with antibody. These tissue data correlate with the ^29^Si MRI studies and confirmed that enhanced ^29^Si MRI signal arose from tumor sites.

While the current study administered the silicon particles to mice via enema, future emphasis will consider delivery options such as an ingestible slurry for application to other gastrointestinal diseases such as colitis, Crohn disease, esophageal cancer, Barrett esophagus, and stomach cancer. Future experiments will incorporate a washout step to remove unbound particles and enhance the signal-to-noise ratio of the imaging procedure.^59^ While the results presented here represent a significant advance in the development of targeted molecular imaging with silicon particles, future efforts will focus on transitioning from microscale to nanoscale silicon. Ongoing efforts ^13,14^ are underway to improve hyperpolarized ^29^Si signal in nanoscale silicon particles for their development as targeted molecular imaging agents. The simple surface chemistry of silicon particles should allow their development as multiplexed theranostic agents, where different targeting agents and therapeutic drugs can be coupled to the particle surface to detect and treat a variety of disease systems in real time. Continuing improvements to imaging sequence parameters should further improve spatial resolution and allow multi-slice imaging of hyperpolarized ^29^Si particles *in vivo* while facilitating dynamic tracking of particles. With ongoing developments of these methods, targeted ^29^Si particle MRI may further supplement diagnostic imaging of CRC.

This work demonstrated the utility of hyperpolarized silicon particles as targeted molecular imaging agents, with the ability to distinguish MUC1-expressing CRC tumors from chemical and biological controls, with confirmatory tissue immunohistochemistry. Confirmation that antibody-functionalization of the silicon particles does not interfere with the hyperpolarization process, and that the harsh conditions of DNP do not affect the targeting efficacy of the antibody, bode well for future development of hyperpolarized silicon particles in noninvasive disease surveillance. The long-lasting ^29^Si signal in silicon particles allows molecular imaging over the course of more than 10 min, which is significantly longer than most other hyperpolarized contrast agents *in vivo* and allows the particles time for targeted binding with tumor surfaces. The biocompatibility and simple surface chemistry of silicon, along with the non-ionizing and non-invasive deep-tissue imaging of MRI and long-lasting positive contrast from hyperpolarization, make hyperpolarized silicon particles potentially well-suited for targeted molecular imaging.

## Materials and Methods

### Silicon particles

The commercially sourced polydispersible silicon powder (Alfa Aesar) consisted of an average particle size of 2 μm (polycrystalline/amorphous) with 99.9985% purity (natural abundance of ^29^Si, ∼4.7%). The particles (similar to those previously characterized in ^9,15,16^) were analyzed using transmission electron microscopy (TEM) and electron spin resonance spectroscopy (Fig. S2). To determine the size of the particles from TEM images, images were manually segmented and the resultant images processed with CellProfiler v 3.1.5. Images were inverted using module ImageMath to make particles appear white on a black background. Individual particles were detected with the module FindPrimaryObjects using the thresholding strategy ‘Global’, method ‘Minimum cross entropy’, lower bound of 0.7, and no distinguishing of clumped objects. All particles smaller than 30 pixels were ignored. Module MeasureObjectSizeShape then measured the longest diameter of each particle, which was converted from pixels to µm on the basis of the size of the scale bar.

### Particle functionalization

Three types of surface chemistry were used in this study: (1) bare (no changes to the surface); (2) PEGylated; and (3) conjugated with PEG-MUC1 antibody (*214D4*, Millipore). The surface-functionalized particles were formulated as follows: after particle sonication, a primary amine group was added to the particle surface using (3-aminopropyl)triethoxysilane (APTES, Sigma-Aldrich). These primary surface amines then were reacted with the NHS-ester side of the heterobifunctional linker NHS ester-(PEG)_8_-maleimide (SM(PEG)_8_ (Thermo Fisher Scientific), resulting in the PEGylated particles (adapted from Salvati et al ^60^). For the *214D4* antibody particles, the antibody first was reduced using *tris*(2-carboxyethyl)phosphine (TCEP); then the exposed thiol groups were linked to the exposed maleimide groups of the PEGylated particles. In between steps, the particles were subjected to centrifugation and washed 2-3X with buffer. Additional details of these reactions are available in Ref. ^61^, as well as Fig. S3; the figure *Inset* shows a schematic of the functionalized particles.

### 29Si DNP

Silicon samples were placed into small polytetrafluoroethylene sample tubes (∼3 mm internal diameter × 2 cm length) and inserted into the laboratory-constructed DNP device. The sample used in the experiment shown in Fig. 1 consisted of dry particles that were packed into the sample tube (and were later removed from the tube and suspended in buffer directly in front of the MRI scanner, then administered to the mouse). The rest of the samples consisted of a solution of silicon particles suspended in buffer (∼130 μL) and pipetted into the sample tubes, which are microwave invisible and withstand cryogenic exposure. The solid-state DNP device consists of a ∼2.9 T ∼100 mW Gunn diode is frequency-modulated from 80.83-80.90 GHz to transfer polarization from the electrons to the ^29^Si nuclear spins, and is directed to the sample via waveguide and slot antenna. The silicon particles typically reached a nuclear spin polarization value on the order of 1%. Following polarization (∼ 16-18 h), the sample tube was removed and manually warmed to room temperature while being transported to the MRI suite for spectroscopy or imaging studies (*T*_transport_ <1 min). Additional details on the polarizer configuration can be found in References ^15,16^.

### In vitro studies

HuMUC1-expressing human colon cancer cell line HT29-MTX-E12 and non-MUC1 expressing bone marrow stromal cell line HS-5 were seeded at around 50,000 cells per well in 8-well chamber slides (Lab-Tek) and cultured for 72 h. Functionalized silicon microparticles were prepared in medium (100 µg silicon microparticles/mL medium). The particles were subjected to vigorous rotary mechanical agitation, sonicated, and mechanically agitated before they were added to each well (200 µL). The cells were then placed in a humidified chamber with an air:CO_2_ ratio of 95:5 (v/v) at 37°C for 60 min on a cell culture rocker at 10 cycles per min. Following particle incubation, the silicon particle suspension was gently aspirated. Cells were gently washed three times with PBS (400 µL per well) and left in the same buffer for imaging on a microscope (Nikon Eclipse TE300 inverted microscope); the imaging data were processed by NIS Elements software (both, Nikon Instruments).

### Modified ELISA for quantification of 214D4-microparticle functionality

PEGylated microparticles, *214D4*-conjugated pre-DNP microparticles, and *214D4*-conjugated post-DNP microparticles were centrifuged at 16,000*g* for 5 min and washed once with 750 µL PBS. Particles then were blocked with 200 µg/mL bovine serum albumin (BSA) for 30 min, shaking at room temperature. Particles then were washed twice and incubated with 15 µg recombinant biotinylated MUC1 ectodomain (additional details included in Supplementary Materials and Methods) in 200 µL PBS for 1 h, shaking at room temperature. Particles then were washed 3X and incubated in a 1:1,000 dilution of Pierce High Sensitivity Streptavidin-HRP (Thermo Fisher Scientific). Next, particles were washed 3X with PBS and transferred to a 96-well microplate in a total volume of 50 µL PBS per sample. To this was added 100 µL of 1-Step Ultra TMB ELISA substrate (Thermo Fisher Scientific). After 5 min of incubation at room temperature, 100 µL of 2 M H_2_SO_4_ was added to each well to quench the reaction. Reactions then were centrifuged at 16,000*g* for 5 min and supernatants recovered to the 96-well plate. Particle pellets were washed twice with PBS, resuspended in 200 µL PBS, and added to the 96-well plate. Absorbance at 450 nm was then measured for the developed TMB substrate and particles. Data were normalized to particle concentration based on a standard curve of 0 to 2 mg/mL of unfunctionalized particles and then normalized to PEGylated microparticle signal.

### 29Si MR spectroscopy

All imaging and spectroscopy experiments utilized a 7T small animal MR scanner (horizontal bore) located adjacent to the silicon DNP device, as well as a dual-tuned ^1^H/^29^Si litz coil (35 mm ID; Doty Scientific). A small (10 mL) vial of silicon oil was used for sequence calibration/optimization; ^29^Si nuclear spin polarization values were typically on the order of ∼1%. ^29^Si MR spectroscopy was conducted at room temperature using a simple pulse/acquire sequence (α=10°) every 5 min following placement of the sample tube into the scanner; reported *T*_1_ value (∼19 min) did not correct for the slight magnetization loss from each signal acquisition. The spectra were analyzed with TopSpin software (Bruker); following Fourier transform, zero-filling, line broadening (30 Hz), and baseline and phase corrections, the integrated values of the ^29^Si peak were recorded.

### Mouse models

The mouse models used in this study were as follows: (1) normal nude mice; (2) nude mice with a subcutaneous injection of huMUC1-expressing HT29-MTX-E12 cells; (3) huMUC1-expressing *Apc^min/+^* mice with a CRC tumor; (4) *Apc^min/+^*mice with a CRC tumor that does not express huMUC1; and (5) an un-related inducible CRC mouse model known as iKAP ^62^ (inducible KRAS, *Apc* and *p53*). To generate subcutaneous huMUC1-expressing tumors, mice received a single injection of ∼4-6 million HT29-MTX-E12 cells into their flank. Once the tumors grew to sufficient size (>5 mm), the mice were used for the imaging study, where the hyperpolarized particles were directly injected into the tumor.

To generate spontaneous huMUC1-expressing CRC tumors, hemizygous huMUC1-expressing female transgenic mice (JAX #024631) were bred to *Apc^min/+^* male mice (JAX #002020), both on the C57BL/6J background. Local breeding colonies were established from the mice, originally obtained from Jackson Laboratory. To encourage tumor formation in MUC1-positive, *Apc* wild-type mice, 1.6% (w/v) dextran sodium sulfate was added to the drinking water for four 21-day cycles of 7 days on, 14 days off, beginning at 8 weeks of age. Mice receiving dextran sodium sulfate were monitored for signs of distress and tumor formation. Mice of both sexes were monitored for CRC development every two weeks beginning at eight weeks of age using a Storz veterinary endoscope. Briefly, the mice were anesthetized with 5% (v/v) isoflurane gas in an induction chamber and maintained on 2% (v/v) isoflurane throughout the procedure via a nose cone, and body temperature was maintained using veterinary heating pads. The colon was insufflated using PBS administered through the endoscopic sheath. Video was taken of each colonoscopy procedure and videos were archived for visual comparison with MR images; an example colonoscopy video is available in the Supplementary Materials and Methods.

For the ^29^Si MRI studies, the mice were anesthetized with 2% (v/v) isoflurane (in 0.75 L/min O_2_) administered via nose cone on an MR-compatible warming sled. Following DNP, the sample was manually warmed near the front of the MRI scanner, the tube was cut using a ceramic blade, and the particle suspension was collected via syringe. For the mice with a subcutaneous tumor, the particles were directly injected into the tumor volume using a 23-gauge needle. For the mice with an orthotopic tumor (and controls), the particles were administered through the rectum using an 18-gauge soft plastic gavage needle (with an additional 300 μL PBS in the syringe); the base of the plastic gavage needle was wrapped in tape in a tapered fashion to function as a cone plug. Once the particles were administered, the gavage needle was left in place and the syringe was taped to the animal bed to prevent anal leakage of particles. Mice receiving the silicon particles rectally were administered a saline solution enema ∼30 min prior to the imaging study to clear the lower intestinal tract. Following the MR study, the mice were euthanized and the lower intestinal tract (from the rectum to the cecum) was collected, separated into about three pieces, and frozen in OCT for later tissue analysis. After completion of the imaging studies, mouse genotypes were confirmed by a person blinded to the imaging condition. All animal studies were performed in accordance with and approved by The University of Texas MD Anderson Cancer Center Institutional Animal Care and Use Committee (IACUC). All animals used in the experiments were treated humanely in accordance with IACUC guidelines.

### Pre-blocking trial

Fresh 100 mM TCEP solution was generated using 20 mM HEPES buffer (pH 7). The *241D4* antibody at a concentration of 1 mg/mL (150 μL) was incubated with 8 μL of fresh 100 mM TCEP solution for 5-7 min at room temperature. Degassed 1× PBS pH 7 (300 μL) was added to the reaction mixture, and 8.2 μL of Cy5-maleimide (100 mM, Lumiprobe 43080) dissolved in dimethyl sulfoxide was added to the solution. The reaction mixture was kept at 4°C in the dark with constant rotation overnight. The mixture was then purified using a GE Healthcare PD midiTrap G-25 column and eluted off the column using 1000 μL of 1× PBS. The protein and Cy5 concentrations were determined using 280 nm and 649 nm absorbance, respectively. The concentrations of *214D4*-Cy5 conjugate ranged between 0.24-0.16 mg/mL.

Each mouse was subjected to an enema prior to pre-block experiments. Animals were anesthetized and approximately 300 μL of *214D5*-Cy5 was injected via the rectum. The rectum was blocked using an oral gavage syringe, which was removed approximately 5 min after rectum injection. Non-bound antibody was removed from the rectum through peristalsis. Animals were imaged by a procedure similar to that described for other hyperpolarized ^29^Si MRI studies. Hyperpolarized silicon microparticles were injected via the rectum approximately 60 to 90 min post *214D4*-Cy5 injection. After hyperpolarized ^29^Si and ^1^H MRI acquisition, the animals were immediately euthanized and necropsy performed. The colon was flushed with PBS buffer and then re-imaged on IVIS Lumina XR (650 nm [excitation] and 680 nm [emission], Supplementary Materials and Methods). Tumor tissue and cecum from mice were embedded in OCT for further pathologic examination.

### 29Si MRI

MRI experiments utilized the same 7T preclinical scanner and dual-tuned ^1^H/^29^Si coil for co-registered imaging. Immediately following administration of the particles, a ^29^Si MR spectra (α = 10°) was acquired to observe the initial ^29^Si signal level; this information was used to determine the wait time prior to imaging (accounting for the particles’ expected *T*_1_). Typical wait times ranged from 5-20 min. ^29^Si imaging utilized rapid acquisition with refocused echoes (RARE) sequences (α = 90°); ^1^H anatomical imaging used a RARE sequence. All imaging was performed in the coronal plane. Additional details of the imaging sequences for each experiment, as well as image processing protocols, are described in the Supplementary Materials and Methods.

### Immunohistochemistry and histology of mouse colon tissue

After sacrifice, the colon was surgically removed, segmented, and placed in OCT medium (Tissue-Tek). Frozen sections (10-µm thick) were cut using a CM1850 ultraviolet cryostat (Leica Biosystems), mounted onto glass slides, and kept frozen. For histologic studies, sections were stained with hematoxylin and eosin at the MD Anderson Division of Surgery Histology Core facility. Slides were scanned by an Aperio Digital Pathology Slide Scanner with a 20× objective (Leica Biosystems). An Aperio ImageScope was utilized to view and generate images. For immunohistochemistry staining, slides were thawed at room temperature and washed with 1× PBS prior to fixation by 4% (w/v) paraformaldehyde (PFA) in water. After 10 min, the PFA solution was removed and cells were washed in PBS, followed by permeabilization for 10 min with PBS containing 0.2% (v/v) Triton-X. The sections then were blocked in 1% (w/v) BSA in PBS with 0.2% (v/v) Triton-X for 1 h at room temperature, then were incubated with MUC1 ectodomain mouse monoclonal antibody *214D4* at 1:100 dilution or MUC1 cytoplasmic tail rabbit polyclonal antibody CT-1 ^63^ at 1:25 dilution in 1% (w/v) BSA blocking buffer overnight at 4°C.

### Excised mouse intestine

After euthanasia post-MR, colons were excised and flushed gently with PBS; residual particle deposition was visually evaluated and photographed prior to further dissection. After this documentation, the colon was processed for histologic and immunohistochemistry studies as already described.

## Supplementary Materials

Materials and Methods

Fig. S1. Uncropped image from Fig. 1.

Fig. S2. Characterization of silicon microparticles.

Fig. S3. Long-lasting HP ^29^Si MR signal inside tumor volume.

Fig. S4. Photographs of excised mouse intestine following administration of HP silicon particles.

Fig. S5. Brightfield images of *in vitro* assays demonstrating specific binding of MUC1-targeted silicon microparticles pre- and post-DNP.

Fig. S6. Immunostaining of colon tissue sections from the transgenic (human) MUC1-expressing orthotopic CRC mouse models described in the manuscript.

Fig. S7. Comparative statistical analysis of molecular targeting ability of *214D4* antibody functionalized hyperpolarized SiPs.

Fig. S8. Images of *in vivo* blocking experiment from one mouse.

Fig. S9A. HP images and spectroscopy from all mice experiments included in the main text.

Fig. S9B. SNR characterization of ^29^Silicon NMR spectra acquired immediately after injection.

Fig. S10. Hematoxylin and eosin staining of OCT-embedded tumor tissue.

Fig. S11. Recombinant biotinylated MUC1 ectodomain is detected by *214D4* antibody.

Fig. S12. Example of step-by-step image processing for two studies.

Video S1. Multimedia video of mouse colonoscopy to ascertain the presence, number, and location of CRC tumors in the orthotopic mouse models.

## Acknowledgements

The authors would like to thank Ms. L. Bitner and Dr. P.K. Marie (MD Anderson Cancer Center) for experimental assistance, and Drs. S. Kopetz (MD Anderson Cancer Center), M.C. Cassidy (U. of Sydney), and C. Marcus (U. of Copenhagen) for helpful discussions.

## Author contributions

NW, JH, PEC, DC, MCFC and PB conceived of the study and planned the experiments. NW, JH, SP, CVM, JSD, NZM, BJE, JXU, KG, HS and DB carried out the experiments, NW, JH, SP, CVM, SR, JSD, NZM, DGM, SWM, STG, DPW, MCFC, DC, PEC and PB contributed to the interpretation of the results. NW, JH, SP, NZM, MCFC, DC, PEC and PB wrote and edited the manuscript with input from all authors.

## Data and materials availability

All essential data generated or analyzed during this study are included in this published article (and its Supplementary Materials files). Any data not shown are available from the corresponding author on reasonable request. Portions of this work were previously included in thesis submitted to Rice University (JXL, JH, CVM) as part of degree requirements.^59^

