## Supplemental File for "Targeted Molecular MRI of Colorectal Cancer by Antibody Functionalized Hyperpolarized Silicon Particles"

[µ]Undergraduate researcher

**Outline:**

Supplementary Figures 1-11
Supplementary Video 1
Supplementary Section 1-2
Supplementary Figures 12

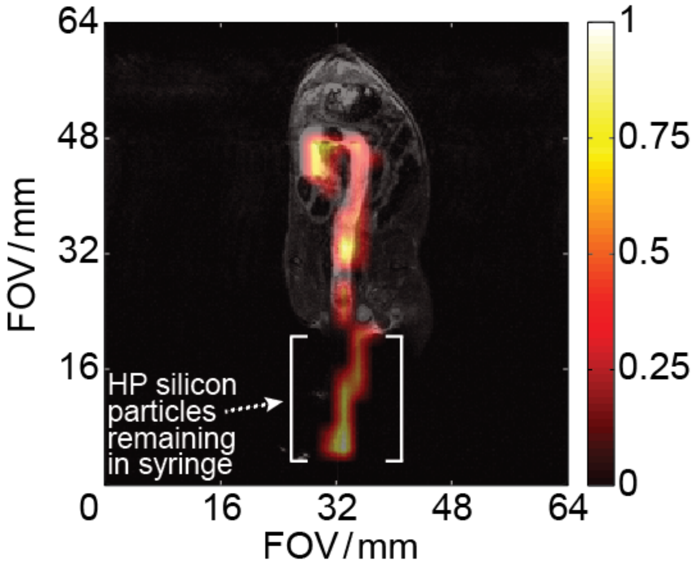


**Supplementary Figure S1:** Uncropped image from Figure 1, displaying the entire field of view. Rectal injection of 125 mg of PEGylated silicon microparticles (in 500 μL PBS) into a wild type mouse; $^{29}$Si image taken 5 minutes post-injection. $^{29}$Si MRI (*color*) overlaid on $^{1}$H MRI (*greyscale*) showing silicon microparticles occupying the intestines from the rectum to the cecum, as well as the HP silicon microparticles remaining in the delivery syringe. Imaging parameters are available in the **Supplemental Section 2**.

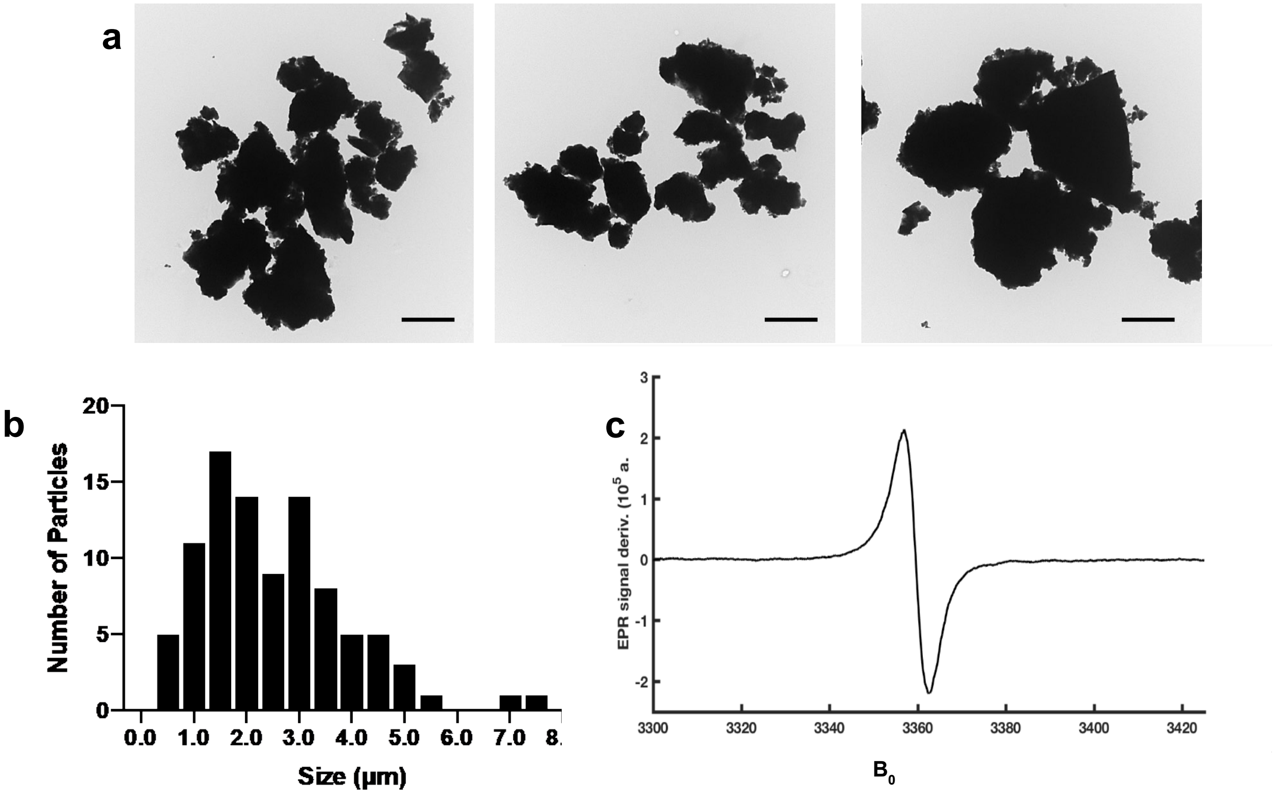


**Supplementary Figure S2:** Characterization of silicon microparticles (SiPs). **a** Transmission Electron Microscope (TEM) images of bare silicon microparticles (scale bar = 2 µm). **b** Size distribution of silicon microparticles based on TEM images of 94 particles. The average size is 2.5 µm. **c** Electron spin resonance (ESR) spectroscopy of the bare particles, showing the presence of endogenous free electrons that can be used for DNP.

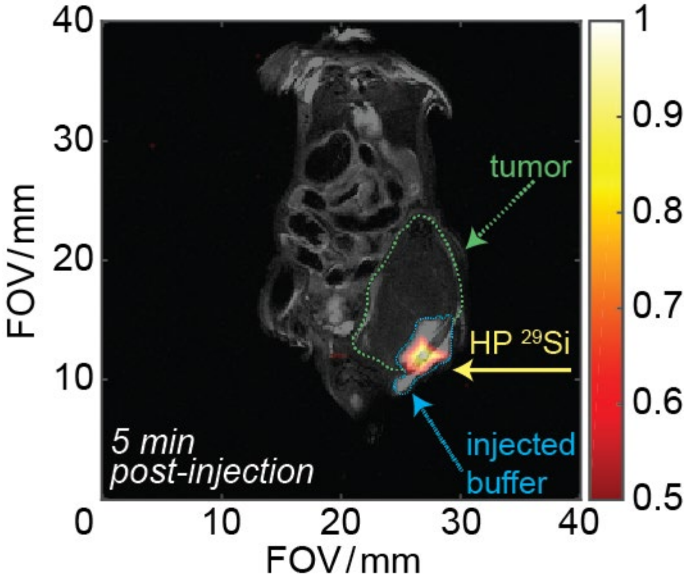


**Supplementary Figure S3:** Long lasting HP $^{29}$Si MR signal inside tumor volume. 60 mg of *214D4*-functionalized 2 μm silicon particles (in 130 μL PBS; $T_{pol}$ ~18 hr) directly injected into tumor volume of a MUC1-expressing subcutaneous HT29-MTX-E12 colorectal cancer mouse model. Image taken 5 minutes after intratumoral injection. Silicon image (*color*) overlaid with $^{1}$H anatomical image (*greyscale*) for co-registration; the tumor volume is outlined with a dotted green line and the $^{1}$H signal from the buffer in the particle solution is outlined with a dotted blue line. Imaging parameters are available in the **Supplemental Section 2**.

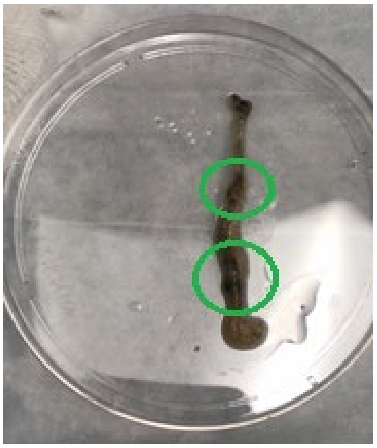

**Supplementary Figure S4:** Photographs of excised mouse intestine following administration of hyperpolarized silicon particles.  Some particles remain after colon excision and gentle flushing with PBS.

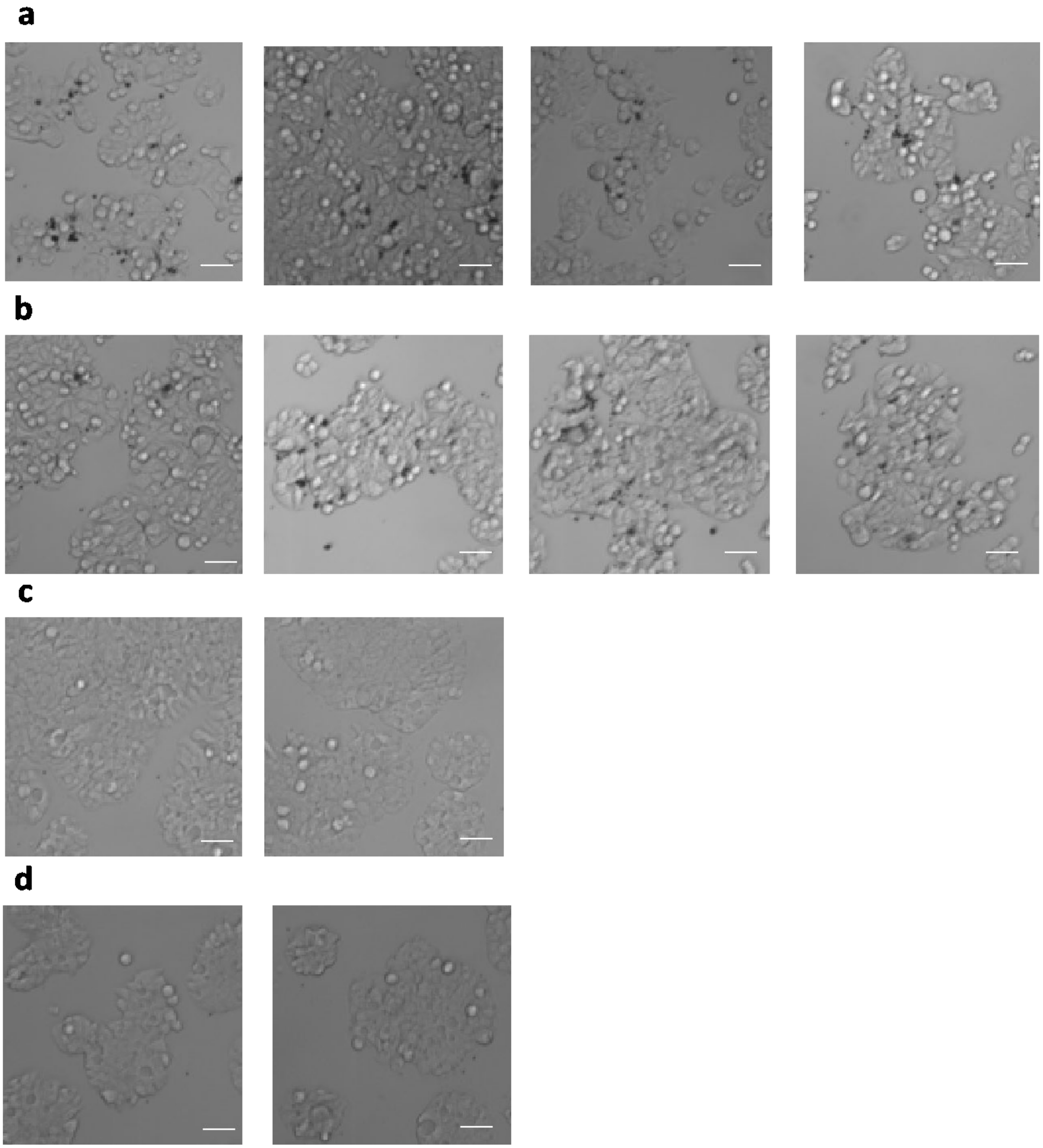
a
b
c
d

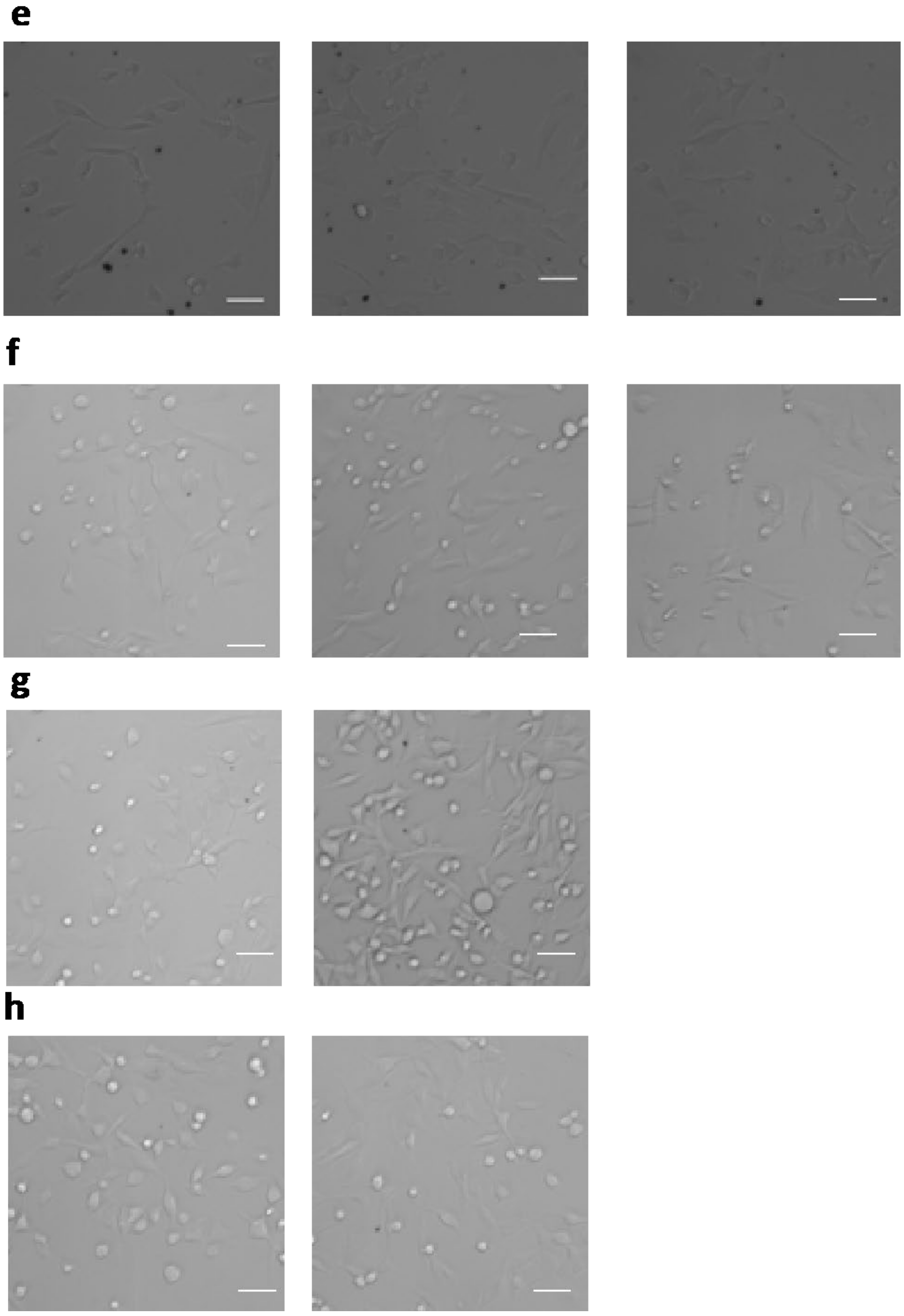

**Supplementary Figure S5:** Brightfield images of *in vitro* assays demonstrating specific binding of MUC-1 targeted silicon micro particles (SiPs) pre- and post- DNP process. **a-d** MUC1 expressing HT29-MTX-E12 cells incubated with SiPs. **a** *214D4*-conjugated SiPs pre-DNP. **b** *214D4*-conjugated SiPs post-DNP. **c** PEG-conjugated SiPs pre-DNP. **d** PEG-conjugated SiPs post-DNP, also referred to as Chemical Control in Figure 3. **e-h** Non-MUC1 expressing HS-5 cells incubated with SiPs. **e** *214D4*-conjugated SiPs pre-DNP.

**f** *214D4*-conjugated SiPs post-DNP, also referred to as Biological Control in Figure 3. **g** PEG-conjugated SiPs pre-DNP. **h** PEG-conjugated SiPs post-DNP. Scale bars indicate 100 μm.

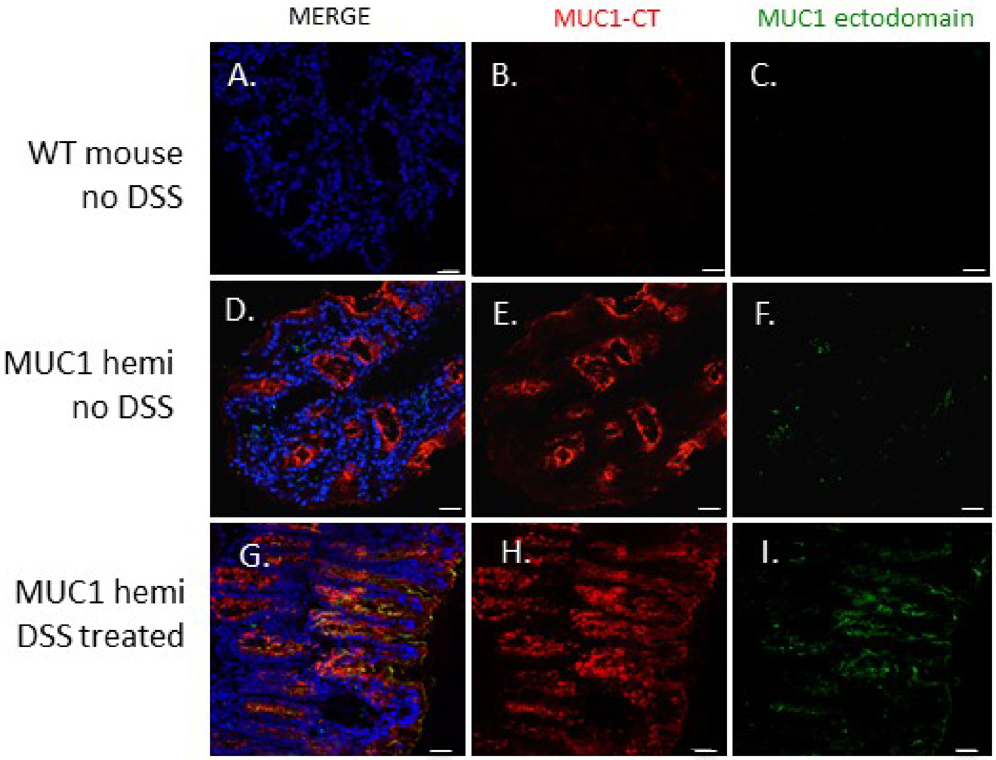


**Supplementary Figure S6:** Immunostaining of colon tissue sections from transgenic (human) MUC1-expressing orthotopic CRC mouse models described in the manuscript. Images show double labeling using CT-1, an antibody that recognizes the cytoplasmic domain of both mouse and human MUC1, and *214D4*, a monoclonal antibody that specifically recognizes the ectodomain of human MUC1. Sections were analyzed by immunofluorescence with anti-CT-1 (red), anti-*214D4* (green), and nuclei (DAPI, blue). Sections from WT mouse colon without DSS treatment (A–C), a human MUC1 hemizygous mouse without DSS treatment (D-F), and a human MUC1 hemizygous mouse with DSS treatment (G-I). Scale bar (lower right) represents 25 µm.

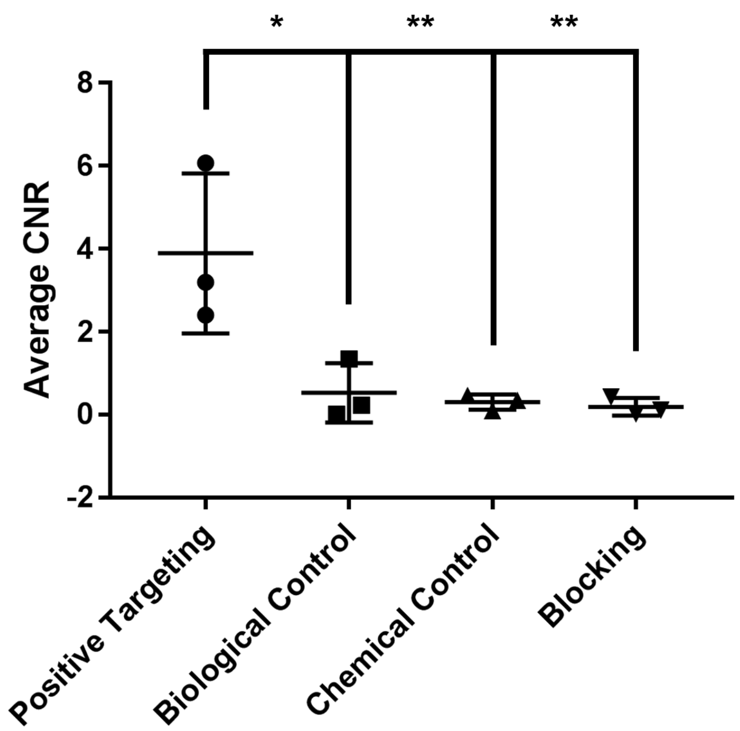


**Supplementary Figure S7:** Comparative statistical analysis of molecular targeting ability of *214D4* antibody functionalized hyperpolarized SiPs through contrast-to-noise ratio (CNR) calculations in MUC1 + animals (N=3), biological (N=3) and chemical controls (N=3) and blocking trials (N=3); * denotes $p<0.05$, ** denotes $p<0.01$.

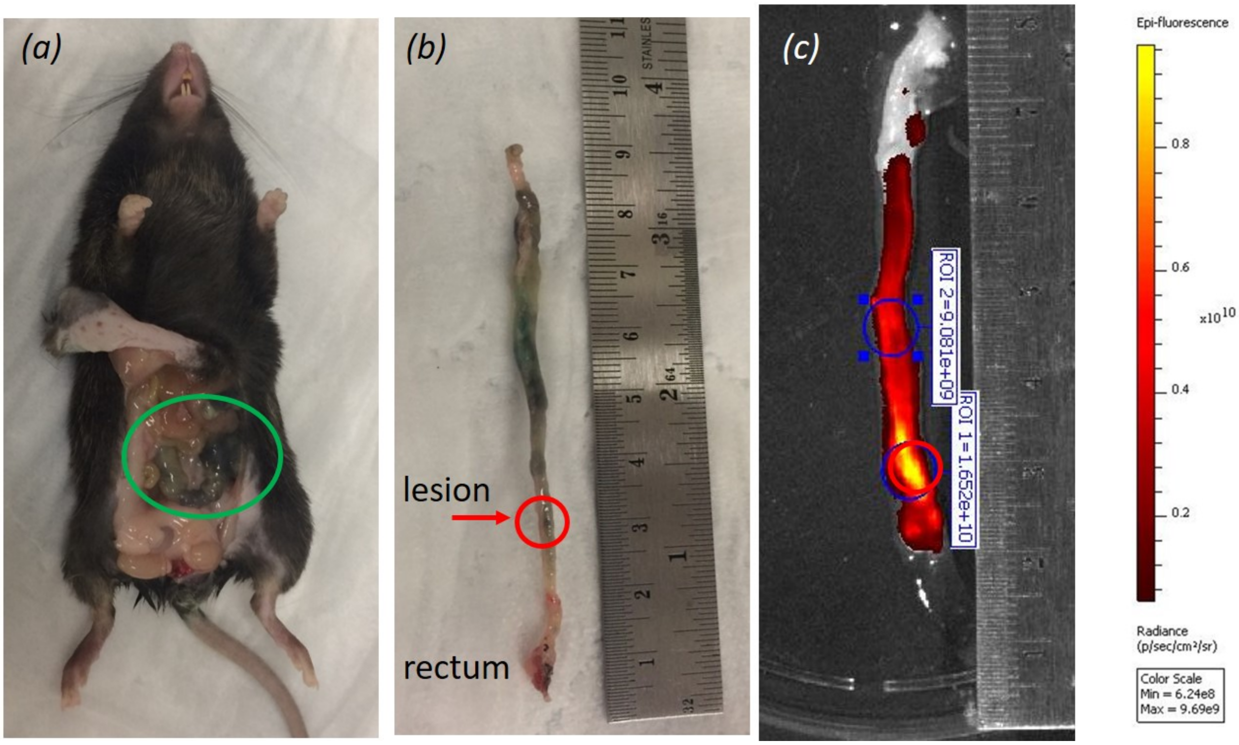


**Supplementary Figure S8:** Images of *in vivo* blocking experiment from one mouse. Experiment was repeated three times. Photographs of excised mouse intestines following administration of 300 ml of *214D4*-Cy5 (blocking) and then hyperpolarized *214D4* conjugated silicon particles: **a** inside mouse, black particle solution can be seen (green circle), **b** some particles and *214D4*-Cy5 remain after colon excision and gentle flushing with PBS. Red circle is the location of a human MUC1-expressing lesion and is in the same location as the area in the IVIS image of the excised colon **c** with the highest Cy5 signal (red circle). In **c** the radiance values for regions of interest (ROIs) are shown outside of the lesion (blue circle) and at the lesion (red circle).

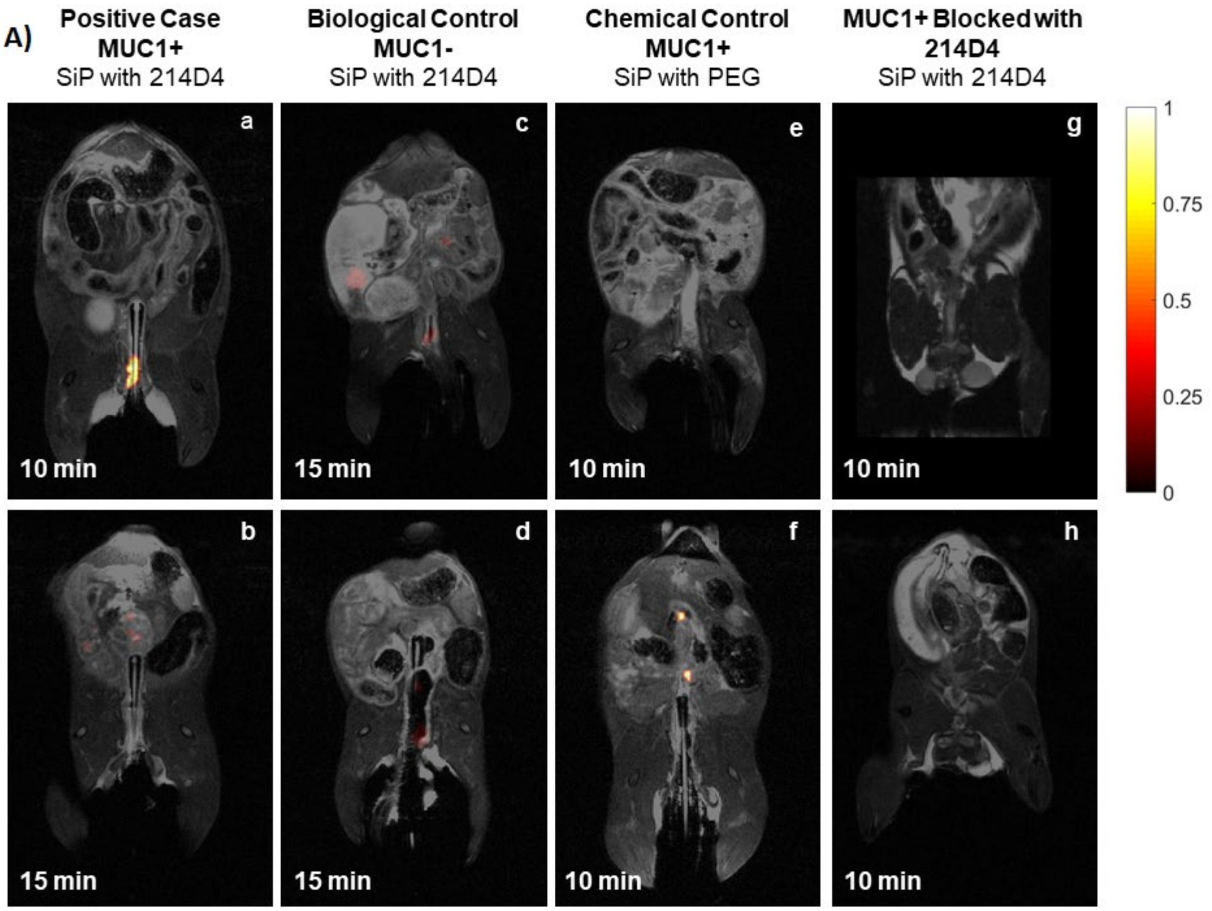


**Supplementary Figure S9A:** Hyperpolarized images and spectroscopy from all mice experiments included in main text. Each image is a separate animal experiment and is an overlay of the proton (Grey) and the hyperpolarized silicon image (yellow to red). HP silicon images were taken 10 to 15 minutes after rectal injection of HP silicon microparticles. The amount of time between injection and image is given. **Panel A) a and b** are images from a mouse expressing human MUC1 CRC and imaged with *214D4* conjugated silicon particles. **c and d** are images from biological controls (mouse possesses colorectal cancer tumors but do not expresses human MUC-1) where *214D4* conjugated silicon particles failed to target the tumor. **e and f** are images from chemical controls (mouse possess colorectal cancer tumor and also expresses MUC-1 but injected with PEGylated particles) where PEGylated particles failed to target tumor. **g and h** Experiments of mouse expressing human MUC1 CRC that was pre-blocked with *214D4*-Cy5 approximately 45 minutes prior to imaging with *214D4* conjugated silicon particles.

**B)**

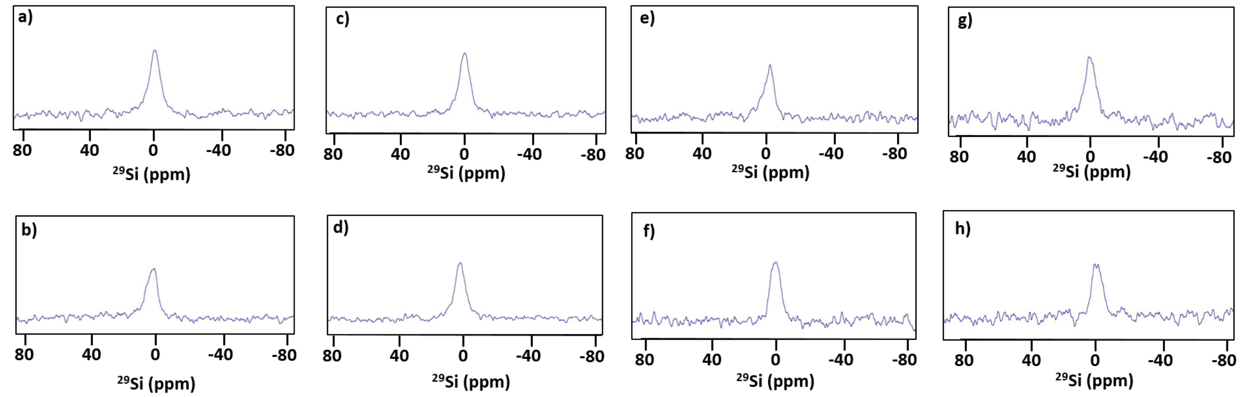


**Supplementary Figure S9B:** To confirm successful initial hyperpolarization and to disprove the trivial explanation for the negative controls, SNR characterization of $^{29}$Silicon NMR spectra acquired immediately after injection are illustrated in the panel-B. It is important to note that $^{29}$Si NMR spectra were taken with $10^0$ pulse immediately after injection of the particles while $^{29}$Si MR images in panel-A (**a to h**) were taken with $90^0$ pulse after 10 minutes post particle injection when the binding event has already taken place. All spectra were acquired with single scan $10^0$ pulse with 2048 time domain points, 101 receiver gain, and spectral width of 100,000 Hz in 7 T horizontal Bruker MR Scanner. All spectra were processed in Topspin 3.1 with zero filling of 4096 points, 60 Hz line broadening, and exponential apodization.

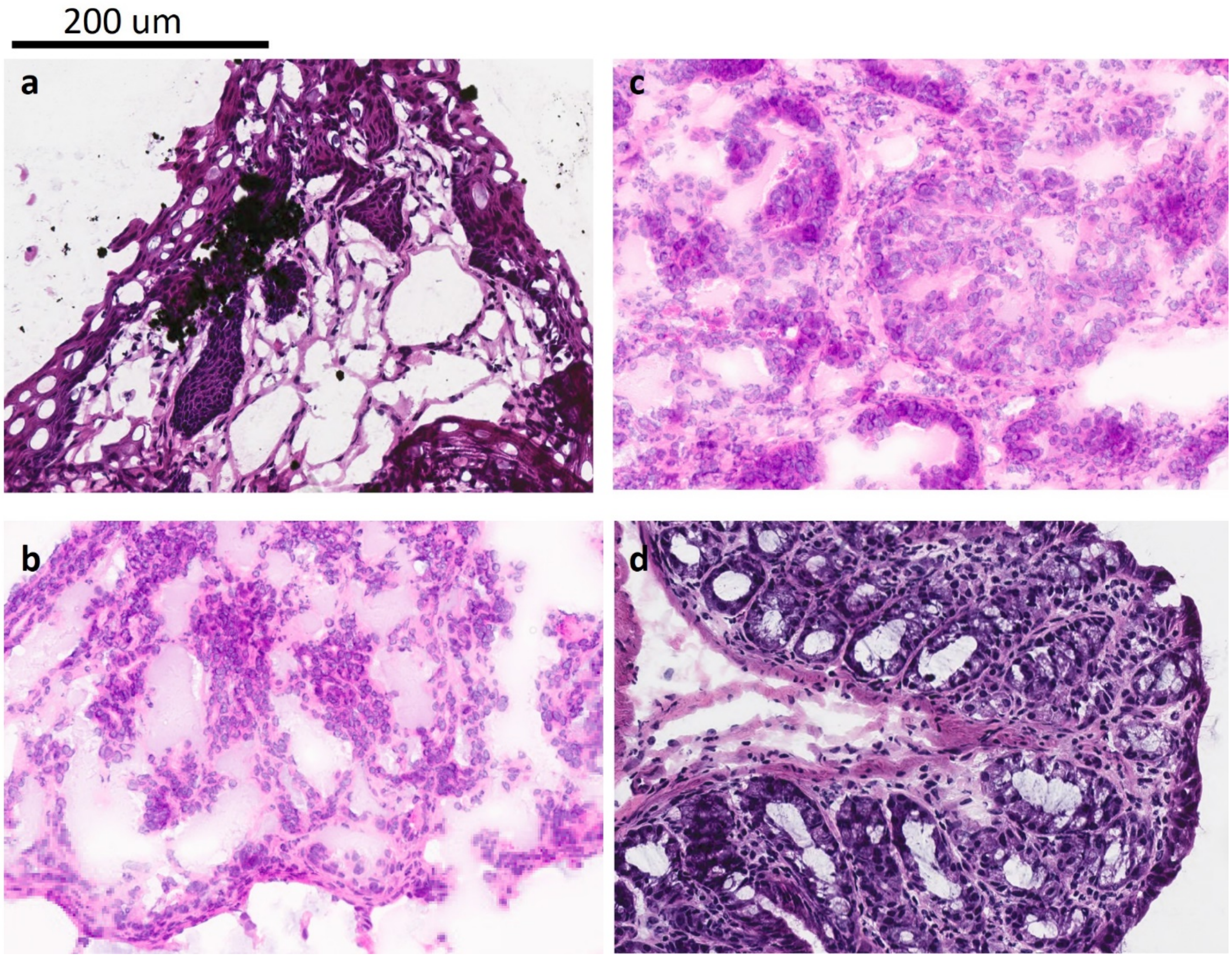


**Supplementary Figure S10:** Haemotoxylin and eosin staining of OCT embedded tumor tissue. **a)** Tissue from mouse expressing human MUC1 CRC and imaged with *214D4* conjugated silicon microparticles (SiPs). Corresponding overlaid proton and silicon image is above in **Supplementary Figure S9a**. **b)** Tissue from mouse expressing mouse MUC1 CRC and imaged with *214D4* conjugated silicon particles. Corresponding overlaid proton and silicon image is **Supplementary Figure S9c**. **c)** Tissue from mouse expressing human MUC1 CRC and imaged with PEGylated silicon particles. Corresponding overlaid proton and silicon image is **Supplementary Figure S9f**. **d)** Tissue from human MUC1 CRC that was pre-blocked with *214D4*-Cy5 approximately 45 minutes prior to imaging with *214D4* conjugated silicon particles. Corresponding overlaid proton and silicon image is **Supplementary Figure S9g**. Black particles are observable only in **a** illustrating the ability of *214D4* silicon particles to target only human MUC1 expressing CRC.

An important point to note that immediately after imaging, the mice were sacrificed and their colons were sectioned and removed. The colon was then gently rinsed with PBS to remove any unbound particles. Post-rinse particle adhesion was only seen in the positive cases. There were control studies that did show measurable signal but because no particles remained after rinsing, we believe this signal originated from aggregation, not specific binding. The signal in control mice was not realized until after image processing, at which point the tissue had already been collected, rinsed, and sectioned. Because the signal in the controls is not apparent real-time, we do not know which parts of the colon should be further studied and looked at histology for what we knew were tumors and anywhere that particles remained after rinsing.

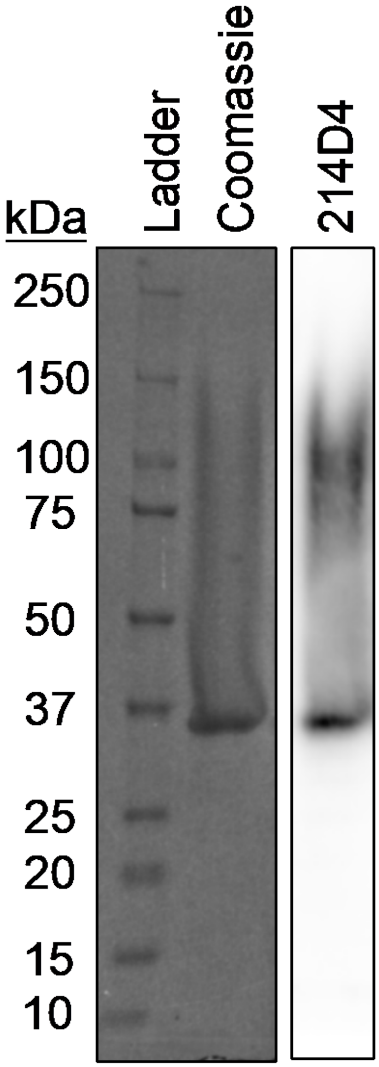


**Supplementary Figure S11:** Recombinant biotinylated MUC1 ectodomain is detected by *214D4* antibody. Coomassie stain (left) and western blot (*214D4*, right) of 2.5 μg of purified protein.

**Supplementary Video SV1:** Multimedia video of mouse colonoscopy to ascertain the presence, number, and location of CRC tumors in the orthotopic mouse models. This procedure was conducted for all orthotopic CRC mice involved in this study. Video files available on request.

Location of the tumor.

Colonoscopy videos were reviewed for tumor appearance and relative location. Specifically, we used the relative distance from the rectum to the point where the colon turns (~4cm from the rectum) to estimate tumor position on the proton image. This distance was further calibrated by the appearance of the gavage tip in the anatomical scan. Upon dissection, the entire colon was removed, flushed gently with PBS and areas with known tumor or apparent particle binding were excised into one OCT block, whereas the remaining colon tissues were harvested into a separate block.

**Supplementary Section 1:**

Functionalization of silicon microparticles with *214D4* antibody.

The following procedure describes how the silicon particles were functionalized with PEG and the *214D4* antibody (adapted from Salvati et al, 2013, *Nat Nanotechnology*, **8**(2): p. 137-43.).

Silicon microparticles were weighed (typical mass ~60 mg), suspended in 70% acidified ethanol (pH 2.5) and sonicated for 5 minutes. (3-aminopropyl)triethoxysilane (APTES, Sigma Aldrich) was added to result in a final concentration of 1.25%, followed by vortex mixing and sonication. The reaction proceeded for ~4 hours on a mechanical rotator at room temperature. Following the reaction, the particles were spun down and washed 3x in 70% ethanol (non-acidic) until a neutral pH was achieved. At this step, a ninhydrin assay *(Rosen, 1957, Arch. Biochem. Biophys*. 67: p. 10–15) was be performed to confirm APTES functionalization by detecting the presence of amine groups. The particles were then re-suspended in 20 mM HEPES buffer (pH 7.4), vortex mixed and sonicated, and washed 2-3x in HEPES buffer. NHS ester-$(PEG)_8$-maleimide ($SM(PEG)_8$) (Themo Fisher Pierce), dissolved in DMSO to a concentration of 240 mM, was added to the particles (suspended in HEPES) at a concentration of 5 mg $SM(PEG)_8$ per mL of HEPES buffer. The particles were vortex mixed and sonicated, and the reaction proceeded using a mechanical rotator at room temperature (at least 4 hours; typically overnight). Following the reaction, the particles were spun down and washed 3x in 20 mM HEPES buffer. This final pellet contained the PEGylated particles.

For the *214D4*-functionalized particles, the antibody solution was added to the PEGylated particles. Preparation of the antibody solution involves: overnight dialysis of 1 mg antibody in 2,000 MWCO dialysis tubing against 2 L of phosphate buffered saline (PBS) at 4 °C to remove Tris (if needed). The dialyzed antibody was transferred into chilled tubes for storage. The reducing agent tris(2-carboxyethyl)phosphine (TCEP) was prepared at a final concentration of 100 mM in $H_2O$ and added to the antibody solution (in HEPES) for a final TCEP concentration of 1 mM; the solution was incubated for ~5 min at room temperature. A desalting column (PD midiTrap—GE Healthcare) was prepared according to the manufacturer's instructions. The reduced antibody solution was run through the column with 20 mM HEPES buffer. The eluation was collected and added to the PEGylated silicon particle suspension (~0.24 mg antibody per 60 mg silicon particles), and allowed to rotate overnight at 4 °C. The particles were then pelleted by centrifugation and washed 2-3x in 20 mM HEPES buffer (pH 7.4) to remove any unreacted antibody. N-acetylcysteine was added to the particle solution (3% w/v) and rotated overnight at 4 °C to form covalent linkages with the maleimide group of any unreacted PEG on the particle surface in order to mitigate particle aggregation. The particles were washed again by resuspension and centrifugation in 20 mM HEPES buffer to remove unreacted cysteine. The final suspension contained 0.1% BSA to prevent particle aggregation and was stored at 4 °C until use (usually within 3 days). An amine-reactive fluorophore can also be added after this step.

Expression, purification and biotinylation of recombinant biotinylated MUC1 ectodomain

A pCDNA3.4 vector encoding ten tandem repeats of the MUC1 ectoodomain (PDTRPAPGSTAPPAHGVTSA x 10), a thrombin cleavage site (LVPRGS), a BirA biotin-protein ligase biotinylation site (GLNDIFEAQKIEWHE) and a hexahistidine tag was ordered from GeneArt (Thermo Fisher Scientific). Protein was expressed using the Expi293F expression system per manufacturer's instructions (Thermo Fisher Scientific). Protein purification was performed at 4 °C with Ni-NTA agarose (G-Biosciences). Briefly, 293F conditioned media was filtered with a 0.2 µM filter and dialyzed 3 x 24 hours against dialysis buffer (25 mM HEPES, pH 8.0, 200 mM NaCl, 5% [v/v] glycerol, 0.01% [w/v] sodium azide). Dialyzed media was mixed 1:1 with equilibration buffer (50 mM phosphate buffer pH 8.0, 300 mM NaCl, 10 mM imidazole, 0.05% [v/v] tween-20). To this was added 1 mL Ni-NTA resin, 20 µL 0.5 M $NiSO_4$ and rotated overnight. Mixture was loaded onto a fritted column and flow-through eluted. Resin was washed once with 25 mL equilibration buffer and twice with 25 mL wash buffer (50 mM phosphate buffer pH 8.0, 300 mM NaCl, 20 mM imidazole). Protein was eluted with 15 mL of elution buffer (50 mM phosphate buffer pH 8.0, 300 mM NaCl, 250 mM imidazole). Eluate was buffer exchanged against dialysis buffer using Amicon Ultracel 10k centrifugal filters (Millipore Sigma). Protein concentration was calculated using the Pierce BCA protein assay (Thermo Fisher Scientific) per manufacturer's instructions. *In vitro* biotinylation of purified protein was performed with recombinant BirA enzyme (Avidity) per manufacturer's instructions.

<u>Western blots</u>

**Cell lysates:** Parental HT29-MTX-E12 and lentiviral transfected GFP- and luciferase-expressing HT29-MTX-E12 cells were separately grown in 6-well tissue culture plates and maintained as described above. At ~90% confluency, protein was extracted from cells with 200 µL sample extraction buffer (SEB) (0.05 M Tris pH 7, 1% (w/v) sodium dodecyl sulfate (SDS), 1% β-mercaptoethanol (BME), 8 M urea, 1:100 dilution phosphatase inhibitor cocktail, and 1:100 dilution phosphatase inhibitor cocktail). Laemmli sample buffer (120 mM Tris-HCl pH 6.8 [Sigma-Aldrich], 20 % (v/v) glycerol [Sigma-Aldrich], 4% (w/v) sodium dodecyl sulfate [SDS; Sigma Aldrich], and 0.02% (v/v) bromophenol blue [Sigma-Aldrich]) was mixed with cell lysate at 1:1 ratio, and incubated at 98 °C for 5 minutes. Protein samples separated by SDS-PAGE with 5% (w/v) polyacrylamide gel and 10% (w/v) polyacrylamide resolving gel at 100 volts for approximately 120 minutes (Laemmli, 1970; Porzio and Pearson, 1977). Protein samples by SDS-PAGE were transferred to nitrocellulose membrane at 40 volts for 5 hours at 4 °C. The transferred blot was blocked with blocking buffer [3% (w/v) bovine serum albumin (BSA) in PBST (PBS with 0.1% (v/v) Tween-20)] for 3 hours at 4 °C, and incubated with primary antibodies *214D4* (Millipore, 1:500 dilution) and □-actin (Abcam 1:500 dilution) overnight at 4 °C. The nitrocellulose blots were washed three times with PBST, 5 minutes each at room temperature, followed with secondary anti-mouse antibody-HRP (1:100,000 dilution) in blocking buffer for 1 hour at room temperature. The blots were washed three times again with PBST, 5 minutes each at room temperature, and developed on audoradiographic film (Denville Scientific) using WestDura ECL (Thermo Scientific) following manufacturer instructions. Finally, ImageJ software was used to perform densitometry on immunoblot images.

**Biotinylated MUC ectodomain:** 2.5 µg of recombinant biotinylated MUC1 ectodomain was boiled in 4x Laemmli Sample Buffer (BioRad) and run on a 4-15% Mini-PROTEAN TGX stain-free precast gel (Bio-Rad Laboratories) at 135V for 50 min. Protein was transferred to a nitrocellulose membrane at 100 V for 1 hour. Membrane then was blocked with 3% BSA (w/v) in TBS containing 0.1% (v/v) tween-20 (blocking buffer) for 1 hour at room temperature. Membrane then was incubated in a 1:2,500 dilution of *214D4* antibody in blocking buffer for 1 hour and washed 3 x 5 min in TBS containing 0.1% (v/v) tween-20. Membrane then was incubated in a 1:10,000 dilution of goat anti-mouse-HRP antibody (Azure Biosystems) in blocking buffer for 1 hour followed by 3 x 5 min washes. Membrane then developed with SuperSignal WestPico substrate (Thermo Fisher Scientific) and imaged with an Azure C600 gel imager (Azure Biosystems).

<u>Immunofluorescence of mouse colon tissue</u>

For immunohistochemistry staining, slides were allowed to thaw at room temperature and washed with 1X PBS prior to fixation by 4% (w/v) paraformaldehyde (PFA) in water. After 10 minutes, PFA solution was removed, and cells were washed in PBS, followed by a 10 minute permeabilization with PBS containing 0.2% (v/v) Triton-X. The sections were then blocked in 1% (w/v) BSA in PBS with 0.2% (v/v) Triton-X for 1 hour at room temperature followed by incubation with MUC1 ectodomain mouse monoclonal antibody *214D4* (Millipore) at 1:100 dilution, or MUC1 cytoplasmic tail rabbit polyclonal antibody CT-1 [61] at 1:25 dilution, in 1% (w/v) BSA blocking buffer overnight at 4 °C. The following day, the primary antibody solution was gently aspirated, and the sections were washed again with PBS with 0.2% (v/v) Triton-X. Secondary anti-mouse AlexaFluor 488 antibody and anti-rabbit AlexaFluor 568 antibody solutions at 1:500 dilution in blocking buffer was added to washed cells, and incubated overnight at 4 °C. The following day, the secondary antibody solution was removed, and the cells were washed with PBS for three times at 5 minutes each. In the last wash, 1 µg/ml: of 4',6-diamidino-2-phenylindole (DAPI) was added for nuclear staining. The solution was aspirated following two minutes of DAPI staining, followed by the addition of Prolong Antifade Mountant solution (Thermo Fisher), and imaged using Nikon A1 Rsi confocal microscope with a 60X water objective.

**Supplementary Section 2:** Sequence parameters and post-processing for MRI scans.

The following describes the imaging and processing parameters for the $^{29}$Si and $^{1}$H MRI scans in individual figures.

**Figure 1:** $^{29}$Si image (*color*): RARE sequence; TR/TE: 59.9 ms/1.8 ms; single average; 64 mm$^2$ FOV with 32 x 32 matrix for a 2 mm$^2$ resolution. Coronal view (single slice). Scan taken 5 minutes after administration of particles (125 mg particles in 500 μL PBS); normal $APC_{min}$ mouse resting posterior. $^{1}$H anatomical image (*greyscale*): RARE sequence; TR/TE: 1926.9 ms/9.5 ms; 3 averages; total scan time: ~3 min; 64 mm$^2$ FOV with 256 x 256 matrix for 0.25 mm$^2$ resolution. Coronal view with 22 slices (0.75 mm slice thickness). Scan taken immediately after $^{29}$Si image was acquired.

**Figures 4 and S3:** $^{29}$Si image (*color*): RARE sequence; TR/TE: 59.9 ms/1.8 ms; single average; 64 mm$^2$ FOV with 32x32 matrix for a 2 mm$^2$ resolution. Coronal view (single slice). Scan taken either 20 min (Fig. 4) or 5 min (Fig. S3) after administration of particles (60 mg particles in 130 μL PBS); $APC_{min}$ mouse with MUC1-expressing subcutaneous tumor, resting anterior. $^{1}$H anatomical image (*greyscale*): RARE sequence; TR/TE: 1,926.9 ms/9.5 ms; 64 mm$^2$ FOV with 256 x 256 matrix for 0.25 mm$^2$ resolution. Coronal view with 22 slices (0.75 mm slice thickness). An anatomical scan was taken immediately after the $^{29}$Si image was acquired. Figure 4 used 4 averages for a total scan time of ~4 min; Figure S3 used 6 averages for a total scan time of ~6 min.

**Figure 5:** $^{29}$Si image (*color*): RARE sequence; TR/TE: 59.9 ms/1.8 ms; single average; 64 mm$^2$ FOV with 32x32 matrix for a 2 mm$^2$ resolution. Coronal view (single slice). Scan taken 10 mins after administration of hyperpolarized silicon particles (45 mg of 2 μm HP SiPs in ~300 μL PBS, time of polarization ($T_{pol}$) ~18 hr, administered through the rectum of mice.); Mouse resting posterior. $^{1}$H anatomical image (*greyscale*): RARE sequence; TR/TE: 1,926.9 ms/9.5 ms; 64 mm$^2$ FOV with 256 x 256 matrix for 0.25 mm$^2$ resolution; 4 averages for a total scan time of ~5 min. Coronal view with 22 slices (0.75 mm slice thickness). An anatomical scan was taken immediately after the $^{29}$Si image was acquired.

**<u>Image Processing and Data Analysis</u>**: All data was processed in Matlab using the following procedure:

a) perform $^{29}Si$ image processing routine:

1. Zero-filling the original k-space data (2D dataset 32x32) to 256x256 before Fourier transformation
2. Import images reconstructed by ParaVision into Matlab
3. Calculate Signal to Noise Ratio (SNR)
   i. Place uniform region of interest (ROI) on tumor locations based on proton image (not on silicon image)
   ii. Calculate average signal intensity (*T*) for tumor ROI
   iii. Place same size region of interest (in the mouse, not machine noise) away from tumor location
   iv. Calculate average signal intensity (*NT*) and standard deviation ($\sigma_{NT}$) for non-tumor ROI
   v. Calculate SNR as follow:

$$\frac{|T - NT|}{\sigma_{NT}}$$

***This is the data that is presented at the Figure 5 bottom-left inset and Supplementary Figure S7.***

4. Zero out signals from left over particles outside the mouse.
5. Apply a threshold to each image to remove signal that is less than five times the background noise, which is calculated by taking the average from 32x32 sections in each corner.
6. Normalize the resulting $^{29}Si$ image
   i. Divide each image by respective NMR data collected from a Silicon oil sample standard to account for variation of the MR scanner
   ii. Divide each image by the largest $^{29}Si$ signal value across all the $^{29}Si$ images and normalize the resulting images into 8bit for display and comparison between studies.

b) perform $^{1}H$ image processing routine:

1. Import images reconstructed by ParaVision (default settings) into Matlab
2. Identify relevant slices and increase the contrast of $^{1}H$ images by saturating the top and bottom 1%

c) overlay the $^{29}Si$ and $^{1}H$ image with the "Image Processing Toolbox" in Matlab.

***This is the data presented in Figure 5 Upper Panel and Supplementary Figure S9A.***

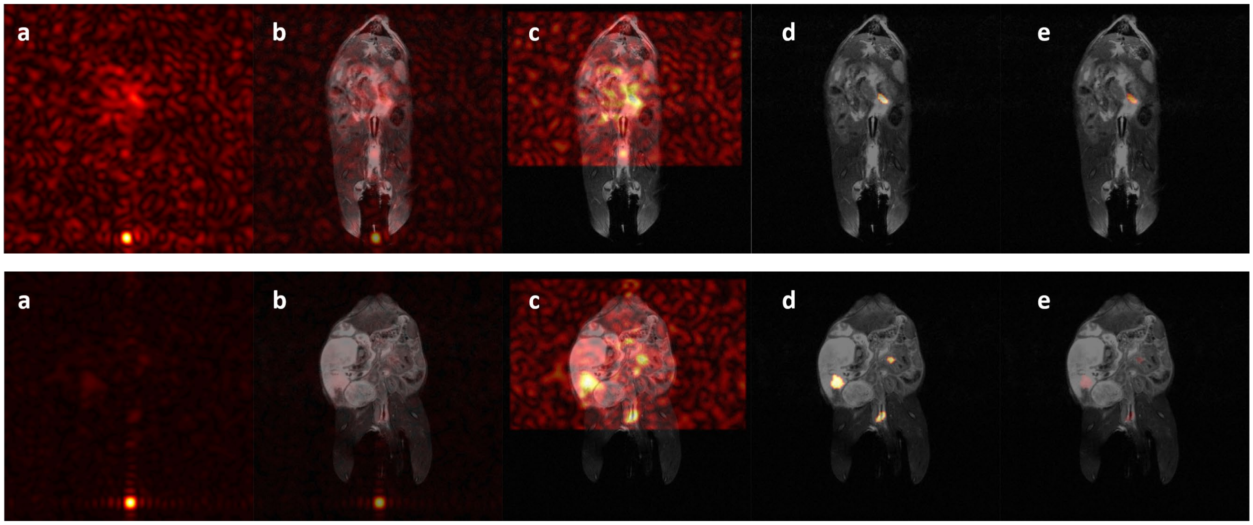


**Supplementary Figure S12:** Example of step by step image processing for two studies. **a)** Raw silicon signal image acquired by hyperpolarized $^{29}Si$ MRI. **b)** Co-registration of silicon signal with anatomical $^{1}H$ MRI. **c)** Signal from particles outside the mouse remaining in the syringe are zeroed out. **d)** Signal less than 5 times the average background noise is zeroed out. Average background noise is calculated from the four corners of the silicon signal. **e)** Silicon signal intensity is normalized to the maximum signal detected across all studies.

Note that thresholding was done between images **c** and **d** in **Supplementary Fig S12**. On the contrary, the quantitative SNR values in the bottom-left inset of **Figure 5** in the main text and the CNR values in the **Supplementary Fig S7** were calculated using the raw silicon image signal without any image processing. No thresholding was done before calculating the quantitative SNR. For this calculation, the SNR was calculated for each mouse independently by comparing the average silicon signal in uniform ROIs centered where tumors were located with same sized ROIs in non-tumor areas of that mouse's gastrointestinal track, all divided by the standard deviation of the non-tumor ROI. ROIs were placed based on the proton image which were then masked to the silicon images for analysis. Tumor locations were determined by MRI and colonoscopy. No image processing was performed before SNR was calculated.
